# The structured mRNA element *45ABC* mediates auto- and cross-regulation of RBP45 genes via alternative splicing

**DOI:** 10.1101/2025.11.25.690383

**Authors:** Maren Reinhardt, Rica Burgardt, Christoph Engel, Maximilian Sack, Zasha Weinberg, Andreas Wachter

## Abstract

Alternative splicing (AS) is a common gene regulatory mechanism involving distinct interactions between *trans*-acting factors and *cis*-regulatory elements on the precursor mRNA (pre-mRNA). In this study, we have functionally characterized the structured motif *45ABC*, which is located in the pre-mRNAs of RNA-binding protein (RBP) 45 genes in many plant species. Our data revealed that this element mediates a negative auto- and cross-regulatory feedback loop via AS of the three *45ABC*-containing RBP45 genes in *Arabidopsis thaliana*. We identified a G-rich stretch within the first stem as a potential RBP45 binding site and observed increased RBP45-dependent AS upon structural weakening of this pairing element. The second stem includes the alternative 5’ splice site being activated in the presence of RBP45. Based on the known interaction between RBP45 homologs and U1 snRNP components required for 5’ splice site recognition, we propose that RBP45 binding to stem I of *45ABC* induces usage of the alternative 5’ splice site in stem II. The resulting splicing variant is unproductive, thereby diminishing RBP45 expression. Analysing the splicing regulatory impact of the three *At*-RBP45 genes in auto- and cross-regulation and a transcriptome-wide manner revealed unequal genetic redundance with a major role of *RBP45B*. Furthermore, phenotypical analysis of single and higher order *rbp45* mutants pointed at these genes’ functions in controlling primary root growth and flowering time. Taken together, we demonstrated that both sequence and structural features of *45ABC* are critical for proper splicing control, balancing RBP45 expression and functions in plants via a conserved mRNA motif.

**Significance statement:** Functional characterisation of a structured mRNA motif present in plant RBP45 genes identified sequence and pairing elements underlying a negative auto- and cross-regulatory expression circuit on the level of alternative splicing. Our study provides a rationale for the evolutionary conservation of this RNA element, which allows balancing levels and functions of RBP45 proteins as a requirement for proper plant development.

## Introduction

During their life cycle, plants need to continuously adapt to environmental fluctuations, involving precise genetic control to fine-tune cellular and developmental processes. A critical layer of this regulation is the splicing of precursor messenger RNAs (pre-mRNAs), a co- or post-transcriptional process that can occur in an either constitutive or alternative manner. While during constitutive splicing the same exon sequences are incorporated in the same order in the mature mRNA, multiple mRNA isoforms are generated by alternative splicing (AS) due to differential 5’ and 3’ splice site selection. For *Arabidopsis thaliana*, an average of 4.4 transcript isoforms per gene was reported to be generated by AS and the usage of alternative transcript start and end sites (Zhang *et al*., 2022). Numerous studies have provided evidence that plants exploit the potential of AS in fine-tuning gene expression to modulate development, responses to stress, and adaptation to environmental changes (Alhabsi *et al*., 2025; Reddy *et al*., 2013; Staiger and Brown, 2013). AS can expand both transcriptomic and proteomic diversity. A substantial proportion of the AS variants is likely not translated into proteins but instead may influence gene expression by altering other processes such as mRNA stability and localization. One widespread mechanism represents the coupling of AS and nonsense-mediated decay (NMD), which causes turnover of non-productive isoforms and thereby allows quantitative gene control (Drechsel *et al*., 2013; Kalyna *et al*., 2012).

Eukaryotic pre-mRNA splicing is regulated by the interplay of *cis*-regulatory elements, specific sequences and structures within the target RNAs, and *trans*-acting factors, including RNA and protein molecules. The splicing regulatory proteins include two major classes: the serine/arginine-rich (SR) proteins and heterogeneous nuclear ribonucleoprotein (hnRNP) proteins. Originally, SR and hnRNP proteins, respectively, have been mainly considered as activators and repressors of splicing sites, however, it is now clear that their action is highly context-dependent and shaped by parameters such as the binding location and the presence of other regulatory factors and elements (Fu and Ares, 2014; Reddy *et al*., 2013). Among the diverse group of hnRNP proteins from *A. thaliana* (Wachter *et al*., 2012), the glycine-rich RNA-binding proteins GRP7 and GRP8 (Streitner *et al*., 2012) and polypyrimidine tract-binding proteins (PTBs; Rühl *et al*., 2012) have been intensively studied and linked to AS. Both GRPs and PTBs are part of auto- and cross-regulatory circuits, triggering formation of NMD-targeted splicing variants from their own and the homologs’ pre-mRNAs when the levels of the respective proteins are elevated. This type of balancing mechanism is common among splicing regulators and other RBPs in plants and other organisms (Müller-McNicoll *et al*., 2019; Reddy *et al*., 2013; Wachter *et al*., 2012).

The RNA-binding protein (RBP) 45 family also belongs to the hnRNP proteins and comprises in *A. thaliana* the four members RBP45A, RBP45B, RBP45C, and RBP45D, each of which contain three RNA recognition motifs (RRMs; Lorković *et al*., 2000). So far, their specific functions and in particular their possible role in AS regulation remain largely unresolved. Previous work has shown that RBP45D interacts with pre-mRNA-processing factor 39a (PRP39a), a component of the U1 small nuclear ribonucleoprotein (snRNP), suggesting a function in 5′ splice site selection (Chang *et al*., 2022). Moreover, RBP45D was found to promote flowering (Wang *et al*., 2022). *In vitro* interaction studies with RBP45B confirmed its RNA binding (Peal *et al*., 2011), and RBP45 proteins were found to associate with proteins involved in various aspects of RNA processing, including U1 snRNP components (Mangilet *et al*., 2024), cap binding protein 20 (CBP20) and the poly(A)-binding protein PAB8 (Muthuramalingam *et al*., 2016), and the RNA helicase up-frameshift 1 (UPF1; Chicois *et al*., 2018; Sulkowska *et al*., 2020). Recently, the presence of an evolutionary conserved structured RNA (strucRNA), *45ABC*, was reported for the pre-mRNAs of *RBP45A*, *RBP45B*, and *RBP45C*, but not *RBP45D* (Sack *et al*., 2025).

In addition to *trans*-acting factors, RNA sequence and structural motifs play a crucial role in facilitating exon and intron definition (Buratti and Baralle, 2004; Georgakopoulos-Soares *et al*., 2022; Ullah *et al*., 2018). For instance, local RNA structures can alter the accessibility of *cis*-regulatory elements, thereby either hindering or enhancing spliceosomal assembly (Shepard and Hertel, 2008). The ability of RNA to form dynamic structures gives rise to a structural complexity that can be responsive to various external factors such as temperature (Chung *et al*., 2020), subcellular localization (Sun *et al*., 2019), and interactions with RBPs (Gosai *et al*., 2015) or metabolite ligands (Steinert *et al*., 2017). RNA secondary structures have been shown to influence diverse steps of gene expression such as AS (Wachter, 2014), RNA localization (Fernández-Moya *et al*., 2021), and RNA stability (Fischer *et al*., 2020; Goodarzi *et al*., 2012). While AS regulation through strucRNAs is well documented in animals (Raker *et al*., 2009; Shepard and Hertel, 2008), few such instances have been reported in plants. One example is the thiamine pyrophosphate (TPP)-binding riboswitch, a conserved RNA structural element found across all three domains of life, including fungi and plants, but being absent from animals (Bocobza *et al*., 2007; Sudarsan *et al*., 2003; Wachter *et al*., 2007). Upon binding of its ligand TPP, the riboswitch’s conformation changes, demasking an alternative splice site and thereby altering the splicing pattern and downregulating the thiamin biosynthesis gene *THIC* in a negative feedback loop. Another example for an RNA structure that is conserved in plants and involved in AS control is the plant 5S ribosomal RNA structural mimic (P5SM; Hammond *et al*., 2009). It interacts with a ribosomal protein to modulate AS of a transcription factor pre-mRNA, adjusting 5S rRNA production to the level of its binding partner L5 (Hammond *et al*., 2009).

In this study, we have characterized the AS-regulatory function of a structured mRNA element that was recently found in a bioinformatic survey of the non-coding regions of plant pre-mRNAs (Sack *et al*., 2025). The strucRNA *45ABC* consists of two hairpins, which are highly conserved among homologs of *RBP45A*, *RBP45B,* and *RBP45C* from monocots and dicots. It consistently overlaps with a cassette exon (CE), the skipping of which generates a splicing variant encoding the full-length RBP45 protein, while its inclusion introduces a premature termination codon and triggers degradation via NMD (Sack et al., 2025). Using splicing reporters based on the three *At*-RBP45 genes, we revealed negative feedback regulation of their pre-mRNAs via AS. Disruption of the motif’s first hairpin resulted in a strong increase in the CE variant, while compensatory mutations restored the authentic splicing output. Further mutational studies identified within this stem I a G-rich sequence that is required for efficient CE inclusion and is proposed to serve as an RBP45 binding site. Moreover, we have established in *A. thaliana* individual overexpression lines as well as single and higher order *rbp45* knockout mutants to examine these genes’ regulatory crosstalk as well as functions in global AS control and plant growth. Transcriptome-wide analyses provided evidence that out of these three RBP45 genes from *A. thaliana* only *RBP45B* plays a broader role in AS control beyond the cross-regulatory circuit. Furthermore, all of the tested loss-of-function mutants exhibited significantly shorter primary roots compared to wild-type plants. In summary, our study has unravelled novel functions of RBP45 genes in *A. thaliana* and identified the strucRNA *45ABC* as a mediator of an AS-based gene regulatory circuit that achieves specificity due to a combination of sequence and structural features.

## Results

### The *45ABC* motif overlaps with alternative splicing sites of *RBP45* genes

Bioinformatic analysis of non-coding regions in available plant genome data revealed several conserved strucRNA candidates that overlap with alternative splice sites (Sack *et al*., 2025). One promising candidate from this study had previously been identified as a conserved intronic element in RBP genes from angiosperms (Burgess and Freeling, 2014), however, its functional role remained unknown. Since it is associated with *RBP45A* (*AT5G54900*)*, RBP45B* (*AT1G11650*), and *RBP45C* (*AT4G27000)* from *A. thaliana*, it was named *45ABC* motif. In the data sets analysed, the motif has 448 matches in genes from 112 mono- and dicotyledonous species that are annotated as *RBP45* homologs or are related to *RBP45* (Supplemental Data Set 1). Interestingly, for all *RBP45* homologs, *45ABC* is positioned in a long intron and encompasses a CE (Figure 1a). Moreover, the two predicted hairpin loops of the motif are in the vicinity of or overlap with the alternative splice sites (Figure 1b), with the alternative 3’ splice site being located ∼10 nt upstream of stem I and the alternative 5’ splice site and the U1 snRNP binding motif (consensus sequence: AG/GUAAGU according to Sheth *et al*. (2006)) being embedded in stem II. Analysis of the *RBP45* transcripts revealed the generation of two major AS variants: Skipping of the CE leads to an isoform (*cd)* that is predicted to encode the full-length protein. CE inclusion introduces a premature termination codon (PTC) and a long 3’ untranslated region (UTR) making this isoform (*nc)* probably unproductive. Accordingly, NMD targeting of the CE inclusion variants from these three *RBP45* genes from *A. thaliana* was experimentally confirmed (Drechsel *et al*., 2013; Sack *et al*., 2025). RT-PCR analysis of *RBP45* splicing in different tissues revealed an overall similar splicing output, with the coding variant being the predominant amplification product in all samples (Figure 1c, Figure S1).

**Figure 1:**
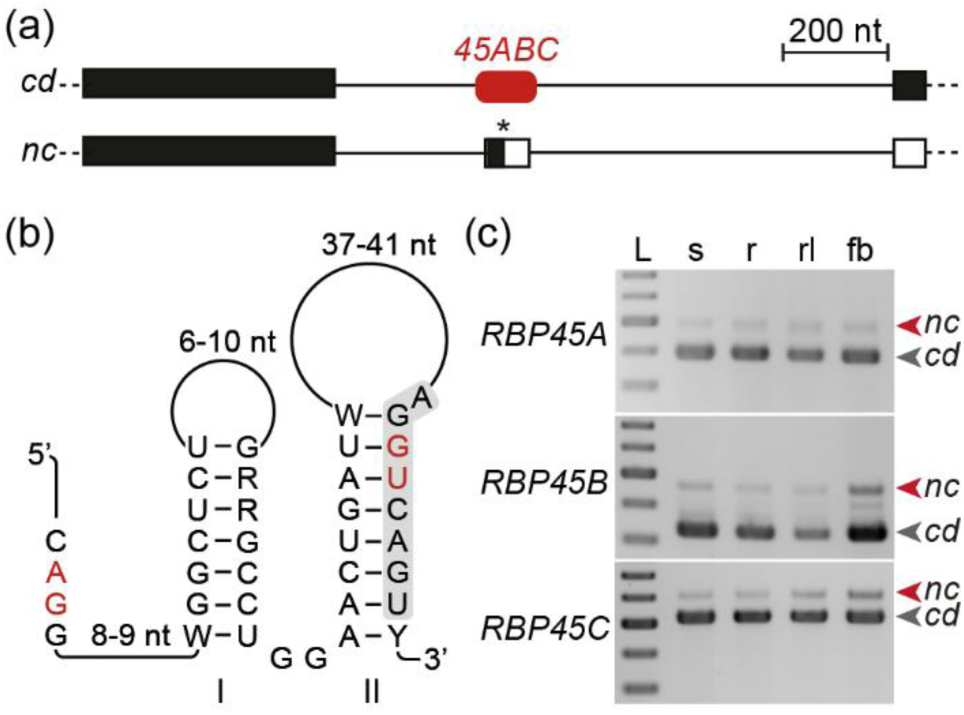
Alternative splicing of *RBP45A*, *RBP45B,* and *RBP45C* genes is associated with the structured element *45ABC*. (a) Schematic depiction of alternatively spliced region of *RBP45A* (*AT5G54900*), *RBP45B* (*AT1G11650*), and *RBP45C* (*AT4G27000*) giving rise to coding (*cd*) and non-coding (*nc*) isoforms with the CE encompassed by the strucRNA *45ABC* (rounded red rectangle). PTC indicated by asterisk; exons, introns, CDS and UTRs are depicted by boxes, lines, black and white shading, respectively. (b) Predicted secondary structure of strucRNA *45ABC* based on the consensus sequence from *A. thaliana*. Red letters show alternative splice sites used for CE inclusion. Letters indicate nucleotides (nt); W, A or U; R, A or G; Y, C or U; lines show variable-length regions. Grey shading marks U1 snRNP binding region. Stems I and II are labelled. (c) AS patterns of *RBP45* genes based on RT-PCR products of samples from 11-day-old whole seedlings (s), roots of 7-day-old seedlings (r), rosette leaves (rl), and flower buds (fb). Binding sites of corresponding primer pairs indicated in Figure S2a. L, size marker in 100 bp increments from 0.3 to 0.7 kb.

### Arabidopsis RBP45 homologues regulate AS of their own pre-mRNAs

The association of the strucRNA *45ABC* with the alternative splice sites in the corresponding pre-mRNAs indicated a functional relationship. To explore a potential regulatory circuit, we designed splicing reporters based on the genomic DNA of *RBP45A, RBP45B*, and *RBP45C* spanning the region from the 5’ UTR to the beginning of the second exon downstream of the CE (Figure 2a). These reporters included an in-frame fusion of the CDS of green fluorescent protein (GFP), resulting in a fluorescent fusion protein upon reporter splicing to the *cd* variant (Figure 2a). Infiltration assays in *Nicotiana benthamiana* leaves were used to examine reporter splicing, in the presence of co-expressed CDS constructs of *RBP45A*, *RBP45B*, or *RBP45C* compared to a control construct containing the CDS of Luciferase (*LUC*). In the *LUC* samples, all reporters showed splicing into both transcript variants, with *cd* being the predominant one (Figure 2b). Co-expression of any of the *RBP45* CDS constructs shifted splicing patterns towards the respective *nc* isoform (Figure 2b, c). This shift in AS towards the *nc* variants correlated with the reduced fluorescence levels of all reporters to less than 50% in the presence of the CDS constructs of *RBP45A*, *RBP45B*, or *RBP45C* (Figure 2d). Taken together, our observations revealed an AS-mediated feedback mechanism in which *RBP45* genes negatively auto- and cross-regulate each other.

**Figure 2:**
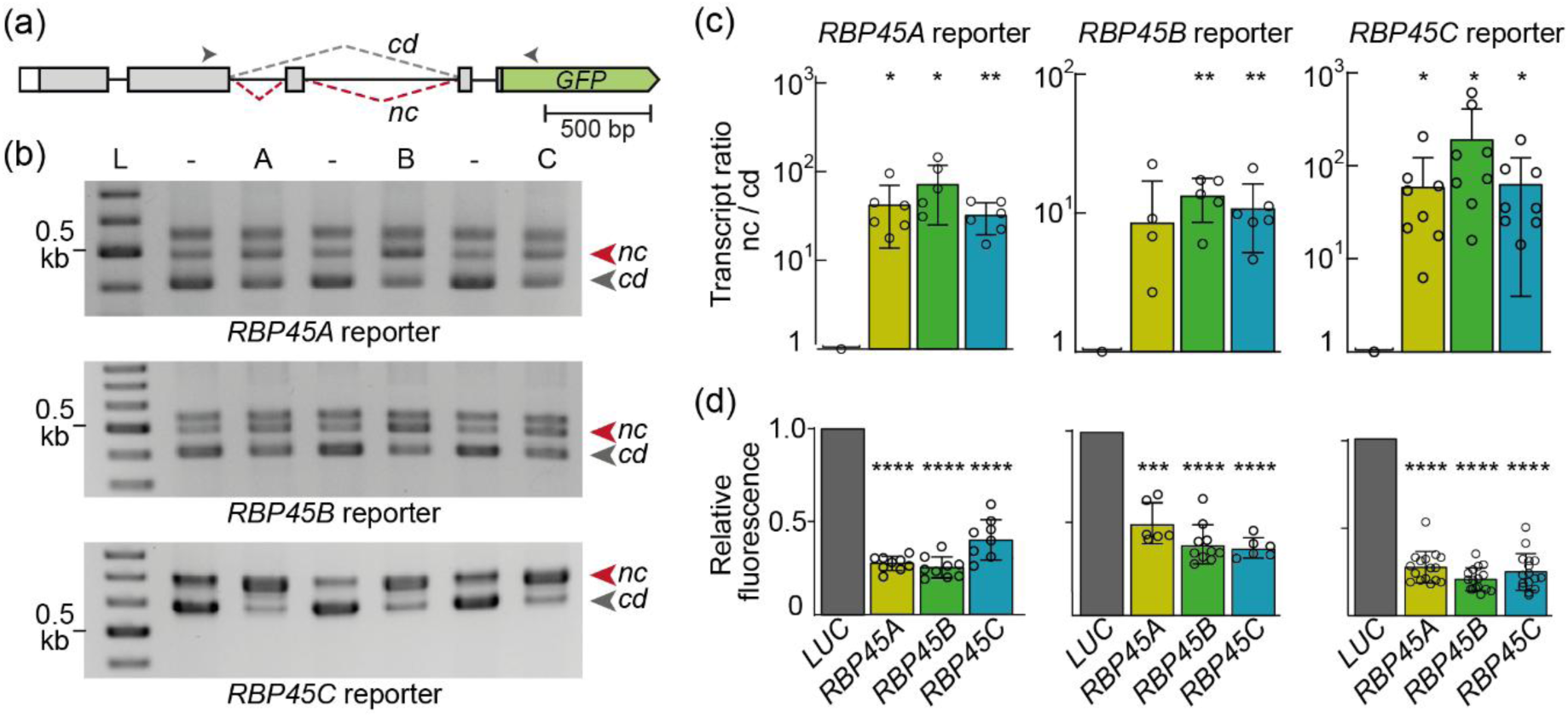
RBP45A, RBP45B, and RBP45C can induce inclusion of the cassette exon overlapping with the *45ABC* strucRNA. (a) Design of splicing reporter constructs for RBP45 genes (display based on *RBP45C)* with boxes and lines representing exons and introns, respectively; arrowheads show the approximate binding sites of primers used for co-amplification PCR shown in (b). (b) RT-PCR products of splicing variants from reporters based on *RBP45A* (top), *RBP45B* (middle), and *RBP45C* (bottom) upon transient expression in *N. benthamiana* leaves co-infiltrated with *LUC* (-), *RBP45A* CDS (A), *RBP45B* CDS (B), or *RBP45C* CDS (C) constructs. The topmost visible band for the *RBP45A* and *RBP45B* reporters likely represents a gel running artefact, as it is absent from Bioanalyzer runs and cannot be identified as distinct splicing variant using sequencing. L: size marker, 100 bp increments. (c, d) Quantification of reporter output based on AS ratios of RT-PCR products analysed via Bioanalyzer (c) and GFP fluorescence (d). Bars indicate mean values, error bars show standard deviations, and open circles represent individual data points. AS ratio and fluorescence of LUC samples each was set to 1. Asterisks indicate significant differences compared to LUC control (one sample t test, *p < 0.05, **p < 0.01, ***p < 0.001, ****p < 0.0001).

To assess this regulatory circuit in its native context, we modulated *RBP45A*, *RBP45B*, and *RBP45C* expression in stably transformed *A. thaliana* plants. This was accomplished either through their overexpression (OE, *35S:RBP45CDS*) or by CRISPR/Cas9-mediated knockout of one, two, or all three corresponding *RBP45* genes (Figure S2). Analysis of total *RBP45* transcript levels confirmed the overaccumulation of *RBP45* mRNAs in the corresponding OE lines (Figure 3a). Notably, the *OE-B* lines showed the strongest extent of overexpression with ∼17 times higher *RBP45* amounts compared to WT. The two independent *OE-A* lines showed an only ∼5-fold overexpression, while for *RBP45C* only one OE line with a less than 2-fold increase in *RBP45C* could be obtained. The other *RBP45C*-OE line probably showed silencing as the level of the coding *RBP45C* transcript from the endogenous locus was decreased and total *RBP45C* levels were WT-like (Figure 3a, b). In general, an increase in the total transcript levels of one of the three *RBP45* genes led to a downregulation of the other two. This effect was particularly pronounced for the paralogs *RBP45A* and *RBP45C.* Furthermore, coding levels of the endogenous genes decreased consistently for all OE lines (Figure 3b). Interestingly, an upregulation of the non-coding transcript variants was only observed in case of the *OE-B* lines, and this effect was consistent among all three genes (Figure 3c), indicating a unique role of RBP45B in alternative splicing control. In contrast, overexpression of *RBP45A* or *RBP45C* resulted in downregulation of the coding and non-coding AS variants from the homologous endogenous genes. This decrease of the non-coding variants in the *RBP45A*-and *RBP45C*-OE lines may be a consequence of their reduced levels of *RBP45B.cd*. Accordingly, the non-coding variants of all three genes may predominantly arise through RBP45B activity in *A. thaliana*. Moreover, the negative auto- and cross-regulation of the *RBP45* genes might involve AS-independent mechanisms.

**Figure 3:**
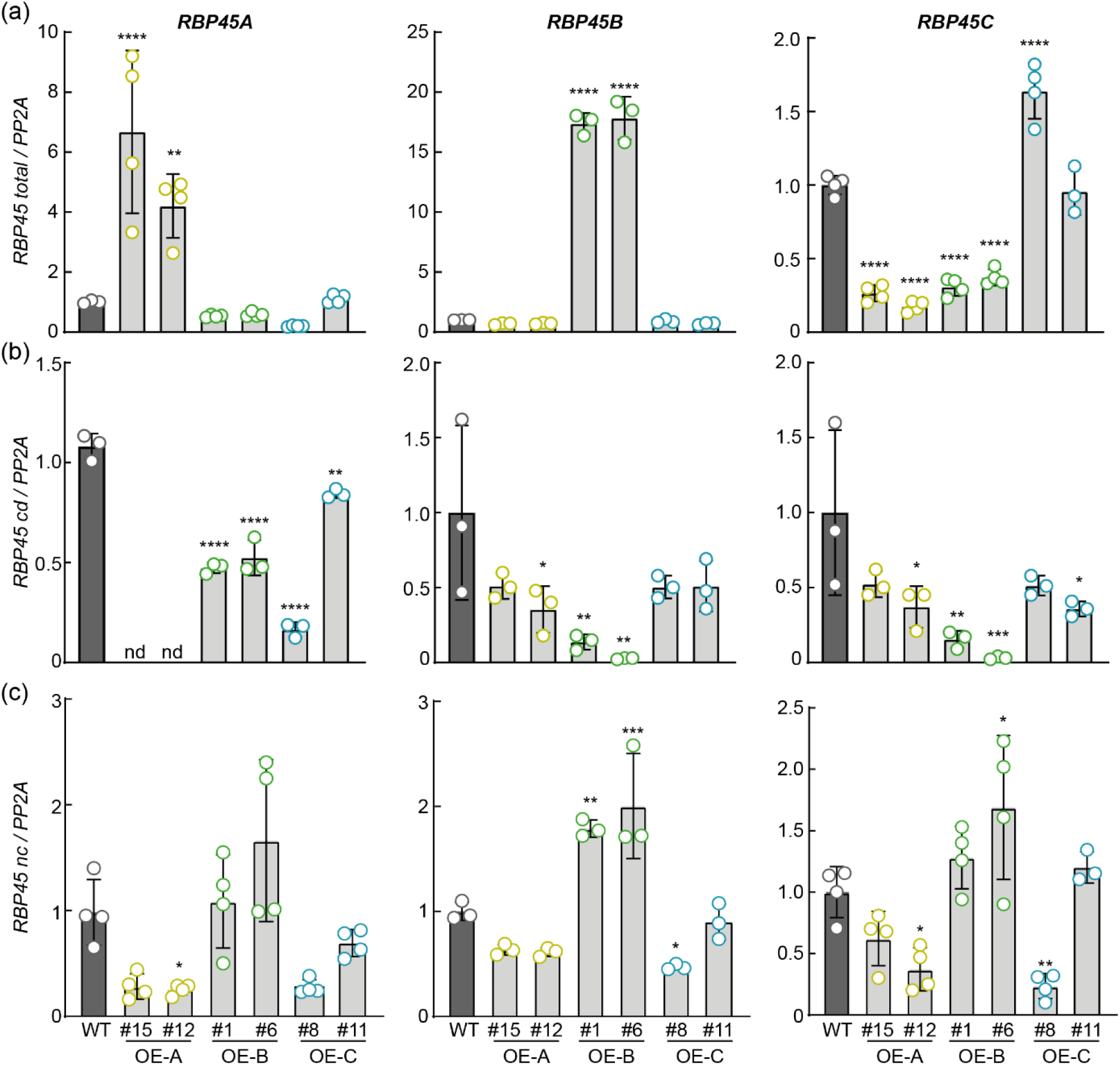
Analysis of negative auto- and cross-regulation in RBP45 overexpression lines unveils a major function of *RBP45B* in splicing control. Quantitative PCR analysis of total (a), endogenous coding (b), and endogenous non-coding (c) RBP45 transcript levels in 11-day-old RBP45 overexpression (*OE*) *A. thaliana* seedlings. All values are expressed relative to reference transcript *PP2A* and normalized to the respective WT mean. Circles represent individual data points from biological replicates and standard deviations are depicted. Asterisks indicate significant change compared to WT (one-way ANOVA followed by Dunnett’s multiple comparisons test, *p < 0.05, **p < 0.01, ***p < 0.001, ****p < 0.0001); nd, not determined.

To complement these findings and further dissect the roles of *RBP45A*, *RBP45B*, and *RBP45C* in AS regulation, we also analysed transcript levels and splicing patterns of the *RBP45* genes in the three mutant lines generated via CRISPR/Cas9 mutagenesis: *rbp45b*, *rbp45bc,* and *rbp45abc* (Figure S2). Given that the mutations give rise to PTCs at early positions within the CDS, the corresponding mutants are considered to be knockout lines. Total transcript levels of the respective mutated genes were significantly reduced compared to those in WT seedlings, and a slight overaccumulation of the non-targeted homolog(s) was seen (Figure 4a). In the *rbp45b* mutant, a decrease of non-coding isoforms was observed for all three *RBP45* genes, whereas the coding transcript variants of *RBP45A* and *RBP45C* showed an opposite trend (Figure 4b, c). This is in line with our conclusion based on the data from the OE lines that *RBP45B* is a major AS regulator for all three *RBP45* genes. Interestingly, knocking out *RBP45C* in addition to *RBP45B* in the double mutant *rbp45bc* restored levels of non-coding *RBP45B* transcript and caused an additional decrease and increase, respectively, of the non-coding and coding *RBP45A* variant (Figure 4b, c). Accordingly, the expression of *RBP45C* and *RBP45A* is tightly linked. Furthermore, the low levels of non-coding variants for all three RBP45 genes in the triple mutant suggested that the cross-regulatory AS circuit might be restricted to this set of genes.

**Figure 4:**
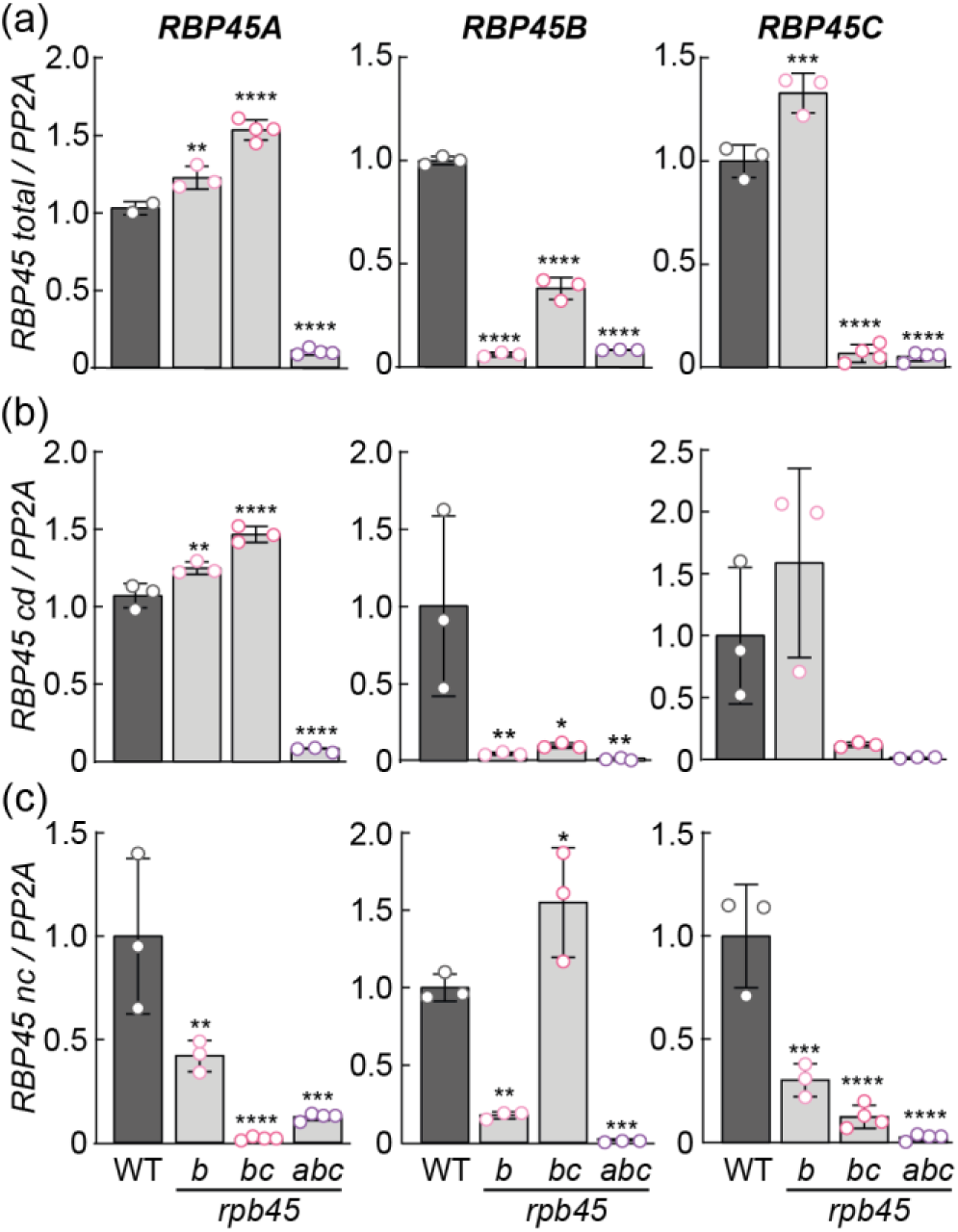
RBP45 knockouts reveal the crosstalk and hierarchy in RBP45-mediated splicing regulation. Quantitative PCR analysis of total (a), coding (b), and non-coding (c) RBP45 transcript levels in 11-day-old *A. thaliana* seedlings in *rbp45* single and higher order mutants. All values are expressed relative to *PP2A* and normalized to the respective WT mean. Circles represent individual data points from biological replicates and standard deviations are depicted. Asterisks indicate significant change compared to WT (one-way ANOVA followed by Dunnett’s multiple comparisons test, * p < 0.05, ** p < 0.01, ***p < 0.001, **** p < 0.0001).

### The structure of *45ABC* is decisive for AS control of RBP45 genes

Our experiments revealed that the three RBP45 genes are subject to negative auto- and cross-regulation via AS. Given the association between the cassette exon and the *45ABC* motif, we next addressed the possible role of this strucRNA in controlling the AS output. To this end, we created the following mutations in the *45ABC* motif in the context of the *RBP45C* splicing reporter (Figure 5a): The disruptive mutants DM1 and DM3 in stem I and stem II, respectively, each involving the exchange of four nucleotides to eliminate base pairing and destabilize the corresponding pairing elements. We ensured that the alternative 5’ splice site within stem II was preserved at its authentic location. To further examine the contribution of the stem I structure, we designed an additional mutant DM2, containing three further nucleotide substitutions and resulting in an almost complete loss of the pairing potential within this part of the element. To ensure that any changes observed for the DMs are due to a change in structure and not the altered sequence, compensatory mutations CM1, CM2, and CM3 were created that are expected to restore the original structure in the context of the new sequence (Figure 5a). Accordingly, if the splicing outcome is primarily dependent on structural features of the mutated regions of the *45ABC* motif, the splicing pattern of the DM and CM constructs, respectively, should be disrupted and restored in comparison to the WT reporter.

**Figure 5:**
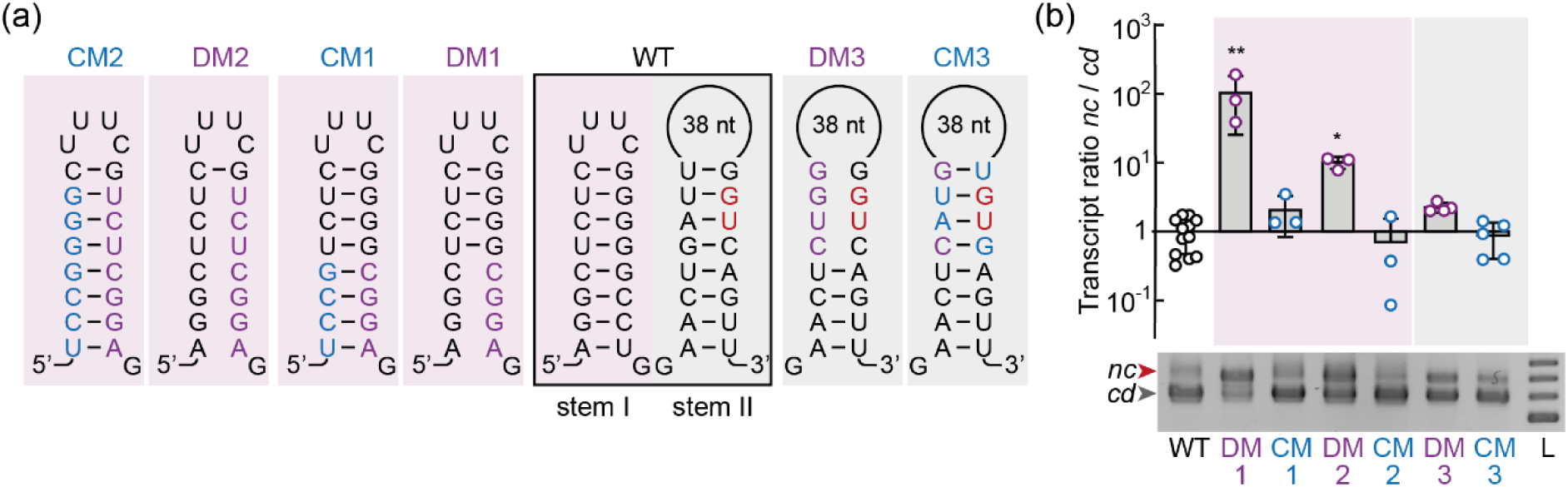
Disrupting the *45ABC* structure causes an AS shift towards cassette exon inclusion. (a) Schematic representation of the base pairing potential of *45ABC* in its wild-type (WT) form of *RBP45C*, as well as disruptive (DM) and compensatory (CM) mutations. Stem I and II areas shaded in pink and grey, respectively. Alternative splice site is indicated in red letters and mutated nucleotides are shown in purple (DM) or blue (CM). (b) Bioanalyzer quantification of WT and mutant *45ABC* reporter AS in 11-day-old stably transformed *A. thaliana* seedlings. Reporter constructs based on *RBP45C* sequence. Upper part shows reporter AS ratios with mean value of WT reporter set to 1. Bars show mean values and individual data points correspond to independent transformant lines. Standard deviations are depicted, and asterisks indicate significant change compared to the WT reporter (one-way ANOVA followed by Dunnett’s multiple comparisons test, * p < 0.05, ** p < 0.01). Lower part shows agarose gel analyses of representative samples with grey and red arrowheads indicating RT-PCR bands from coding and non-coding splice variants, respectively. L: size marker, from bottom to top: 0.5 – 0.8 kb, in 0.1 kb increments.

We initially tested these splicing reporters upon transient expression in *N. benthamiana* leaves and observed that the disruptive mutants DM1 and DM2 in stem I resulted in almost exclusive splicing to the *nc* isoform, even in the absence of RBP45 CDS co-expression (Figure S3a). In case of DM3 in stem II, only a partial splicing shift towards the *nc* variant was seen, and this effect was further enhanced upon RBP45 co-expression. All compensatory mutations reverted this splicing pattern by causing a relative increase of the *cd* variant, however, in comparison to the WT construct, CM1 and CM2 did not fully compensate while CM3 overcompensated. In case of the stem I mutants, the response to RBP45 co-expression was almost completely lost and restored, respectively, for the DM and CM constructs (Figure S3b). As the transient analysis in *N. benthamiana* leaves represents a heterologous system with artificially high expression, we next tested splicing of the various reporter constructs in the endogenous context. In stably transformed *A. thaliana* lines, DM1 led to a significant shift towards the *nc* isoform (Figure 5b). In contrast to the transient system, in the native context CM1 was able to restore the splicing pattern to a WT-like AS ratio. These findings further support that the stem I structure contributes to proper splicing regulation. The more extensive mutation DM2 in stem I also resulted in a shift towards the *nc* isoform, although this effect was quantitatively less pronounced than in case of DM1. Nevertheless, CM2 also fully restored the WT splicing pattern, being in line with the critical role of the stem I structure. In contrast to our observations for the stem I mutants, the stem II-targeting DM3 showed only a slight increase in the *nc*/*cd* ratio. Still, the CM3 mutation reverted this minor splicing shift back to WT levels, indicating that stem II may also contribute to splice site selection (Figure 5b). Collectively, these results demonstrated that the structure of *45ABC*, particularly of stem I, plays a critical role in modulating the AS outcome of the corresponding RBP45 genes.

We hypothesized that the base-pairing within stem I may affect the splicing outcome by controlling the accessibility of an RBP45 binding site. A prior study revealed that recombinantly expressed RBP45 protein from *Nicotiana plumbaginifolia* exhibits *in vitro* binding to poly(U) and poly(G) ribohomopolymers (Lorković *et al*., 2000). Interestingly, a consecutive stretch of guanines is present in stem I of *45ABC* in the case of *RBP45C.* For *RBP45A* and *RBP45B,* one of the guanines is replaced by adenine. Analysis of all available instances of this motif identified in this region the purine-rich sequence RGRG (Sack *et al*., 2025). To experimentally test the relevance of this G-stretch within stem I, additional mutations were introduced into this strucRNA in the context of the *RBP45C* splicing reporter. In the binding mutant BM2, the consecutive guanines were substituted with adenines to assess the role of binding sequence specificity. Moreover, three C nucleobases on the complementary strand were mutated to Us to preserve the pairing potential. As a control, we generated the additional mutant BM1, in which the complementary poly(U) stretch was included while the G-stretch and the pairing potential were preserved (Figure 6a). Upon transient co-expression with the *LUC* control in *N. benthamiana* leaves, BM1 displayed a WT-like splicing pattern, whereas BM2 exhibited a near-complete loss of splicing to the CE-containing variant (Figure 6b, c). This observation provided evidence that the G-stretch is critical for splicing to the *nc* isoform. Upon co-expression with the CDS constructs of any of the three RBP45 homologs, BM2 still showed an AS response, however, the splicing shift was markedly less pronounced compared to the WT and the BM1 mutant. This responsiveness of the BM mutants can also be seen at the protein level of the reporters, with reduced fluorescence upon RBP45 co-expression (Figure S3c). In line with the increased splicing of BM2 to the coding variant, this mutant resulted in higher reporter fluorescence than the WT construct (Figure S3d). The observation that splicing of the BM2 mutant reporter is still responsive to RBP45 co-expression might be explained by massive accumulation of the corresponding RBP45 proteins and rather unspecific RNA binding in the *N. benthamiana* system. Moreover, the introduced U-stretch might also enable RBP45 recruitment given the above-mentioned binding preference observed for a homolog from *N. plumbaginifolia* under *in vitro* conditions (Lorković et al. 2000). Together with the results for the structural mutant reporters (Figure 5 and S3), we propose a combined impact of structure and sequence on AS control. RBP45 binding to the G-rich sequence in stem I is assumed to trigger CE inclusion, e.g. due to structural changes and/or recruitment of further regulators. Disruption of the corresponding pairing element might facilitate RBP45 binding and thereby shift the AS ratio towards more CE inclusion, as observed in DM1 and DM2. The higher number of G residues in the 3’ part of stem I for DM1 compared to DM2 might result in more efficient RBP45 binding, explaining the stronger shift towards the *nc* variant despite less extensive stem weakening in case of DM1.

**Figure 6:**
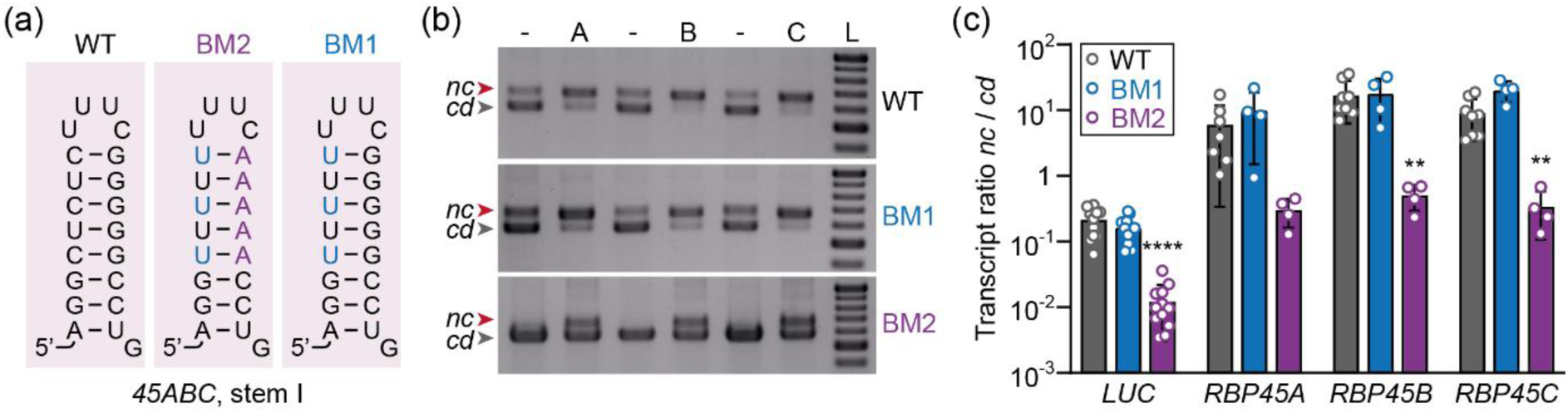
A purine stretch within stem I of *45ABC* promotes inclusion of the cassette exon. (a) Schematic representation of the base pairing potential within stem I of *45ABC* in its wild-type (WT) form as well as potential binding motif mutations (BM1, BM2) in the context of an *RBP45C* splicing reporter. Mutated nucleotides are depicted in blue or purple as in Figure 5a. (b) Agarose gel analysis of RT-PCR products of splicing variants derived from *RBP45C* WT, BM1, and BM2 reporter upon co-expression in *N. benthamiana* leaves with *LUC* (-) or CDS constructs of *RBP45A* (A), *RBP45B* (B), or *RBP45C* (C). L: size marker, from bottom to top: 0.4 – 1.0 kb in 0.1 kb increments. (c) Log-scale quantification of reporter splicing for the samples described in (b). Mean values with standard deviation are depicted; open circles represent individual data points. Asterisks indicate significant differences compared to the WT construct (two-way ANOVA followed by Dunnett’s multiple comparisons test; **p < 0.01, ****p < 0.0001).

### Misexpression of RBP45 can cause reduced primary root length and altered flowering time

To gain insights into the biological relevance of the RBP45 genes, we compared the development of the RBP45 misexpression lines and WT plants. A previous study reported reduced primary root lengths for *rbp45d* mutants, while no such phenotype was seen for *rbp45a* and *rbp45c* T-DNA insertion lines (Huang *et al*., 2022; Muthuramalingam *et al*., 2016). Measuring root lengths of our mutant lines revealed that the loss of *RBP45B* also resulted in shorter roots, and this phenotype was even more pronounced in the corresponding double and triple mutants (Figure 7a). A diminished root length was also observed for the *OE-C* line, whereas the two other overexpression lines showed WT-like growth. To test whether the altered root phenotype of the corresponding lines might be a consequence of diminished seed size, we measured the seed surface area of the mutant lines and different batches of WT seeds. The variation observed among the mutants and different batches of WT seeds were in a similar range (Figure S4). Accordingly, the altered root length can at least not solely be explained by varying seed filling and resource availability. Another developmental parameter affected in some of the mutants was the flowering time. For the knockout mutants, only *rbp45b* showed a slightly delayed onset of flowering (Figure 7b). Later flowering had also been described for *rbp45d* mutants (Huang *et al*., 2022; Muthuramalingam *et al*., 2016), further indicating that RBP45 genes can share common functions in development. Interestingly, the effect of RBP45 overexpression on flowering time differed between the three homologs. While *OE-A* lines showed only a minor delay in flowering onset compared to the WT, *OE-B* and *OE-C* lines, respectively, flowered significantly earlier and later. These findings suggested that RBP45 genes might have specific and redundant regulation targets and that proper expression control of RBP45 genes is critical for normal plant development.

**Figure 7:**
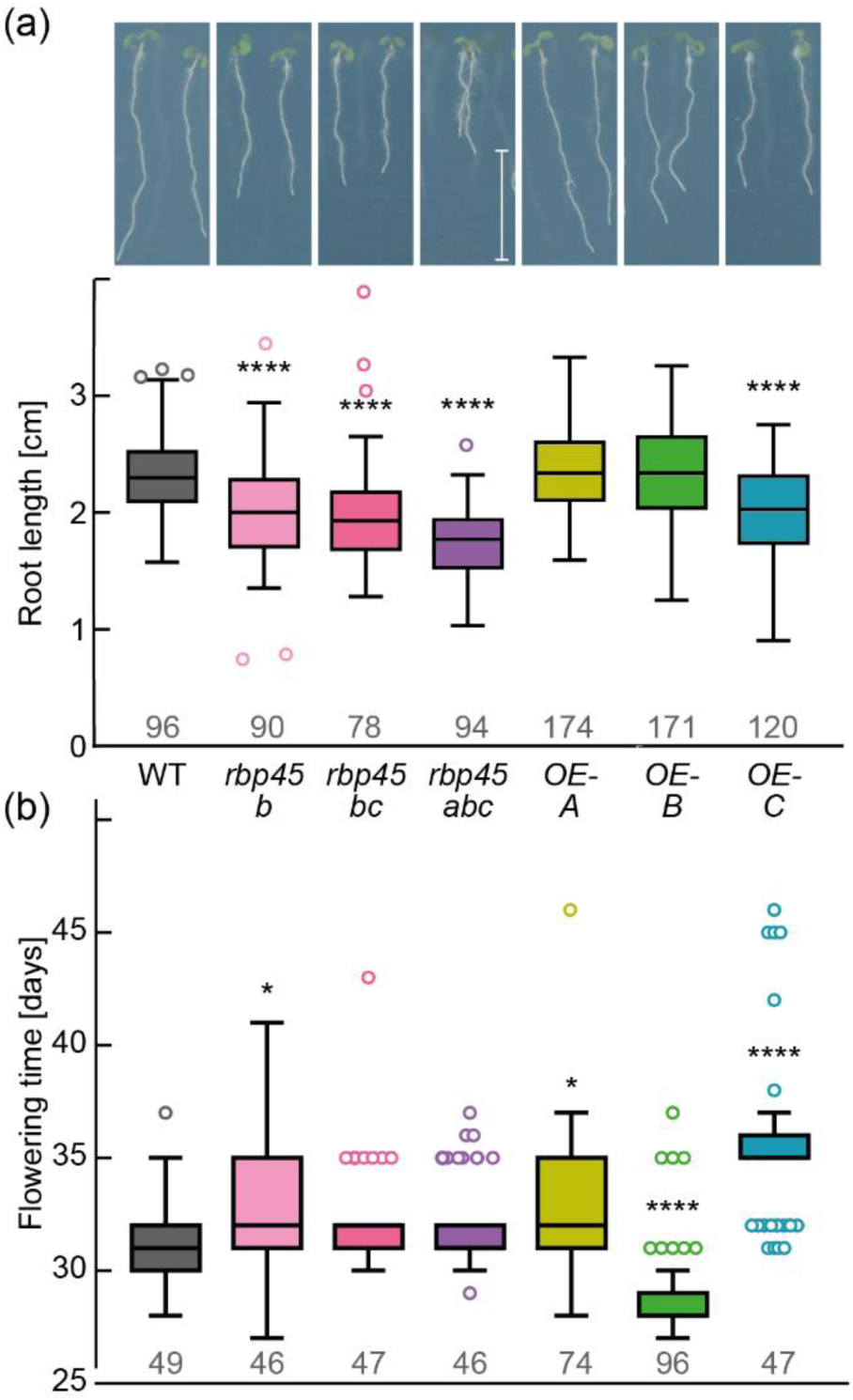
Altered root length and flowering time upon RBP45 misexpression. (a) Primary root lengths of 7-day-old *A. thaliana* WT and RBP45 misexpression lines depicted by representative pictures and mean values with standard deviations. White scale bar corresponds to 1 cm and numbers of analysed seedlings from three independent experiments are indicated. Asterisks indicate significant change compared to WT (one-way ANOVA followed by Dunnett’s multiple comparisons test; **** p < 0.0001). (b) Flowering time in days after sowing. Lines, display details, and statistical analysis (* p < 0.05, **** p < 0.0001) as described for (a).

### RBP45B is the major regulator of AS among the three RBP45 paralogs

Since we observed that the three RBP45 genes are subject to auto- and cross-regulation involving splicing control, we next investigated to which extent the RBP45 genes can influence gene expression and AS at the whole transcriptome level. Therefore, RNA sequencing analyses were performed using *A. thaliana* seedling samples of the higher order mutants *rbp45bc* and *rbp45abc* as well as the constitutive OE lines for *RBP45A*, *RBP45B,* and *RBP45C.* Based on the altered root phenotype, we also performed transcriptome-wide analyses using root tissue of 7-day-old WT seedlings and those mutants displaying shorter primary roots, namely *rbp45bc*, *rbp45abc*, and *OE-C*. Data were analysed via the 3D RNA-seq pipeline (Guo *et al*., 2019), resulting for both datasets in the detection of over 16,000 genes that were expressed in at least one sample type (Figures S5 and S6). The gene expression analysis revealed rather few changes, with 36 and 52 differentially expressed (DE) genes, respectively, in the whole seedlings and root samples (Figure 8a). Only eight of the DE genes were shared between the two datasets, indicating the involvement of tissue-specific regulation. Most of the DE genes were downregulated in the mutants, both for whole seedlings and roots (Figure 8b and Supplemental Data Set 2) and with the largest number of DE genes being detected in *rbp45abc* roots. This observation is in line with the finding that the triple mutant showed the most pronounced reduction in root length. Among the DE genes was a *copia-like retrotransposon* (*AT5G35935*), whose function is uncharacterized yet and which was consistently downregulated in all RBP45 misexpression lines relative to WT samples in both seedlings and roots (Figure S7). Given that the alteration in the transcript level of this retrotransposon showed no correlation with RBP45 expression level, we concluded that this change is probably not directly linked to RBP45 function.

**Figure 8:**
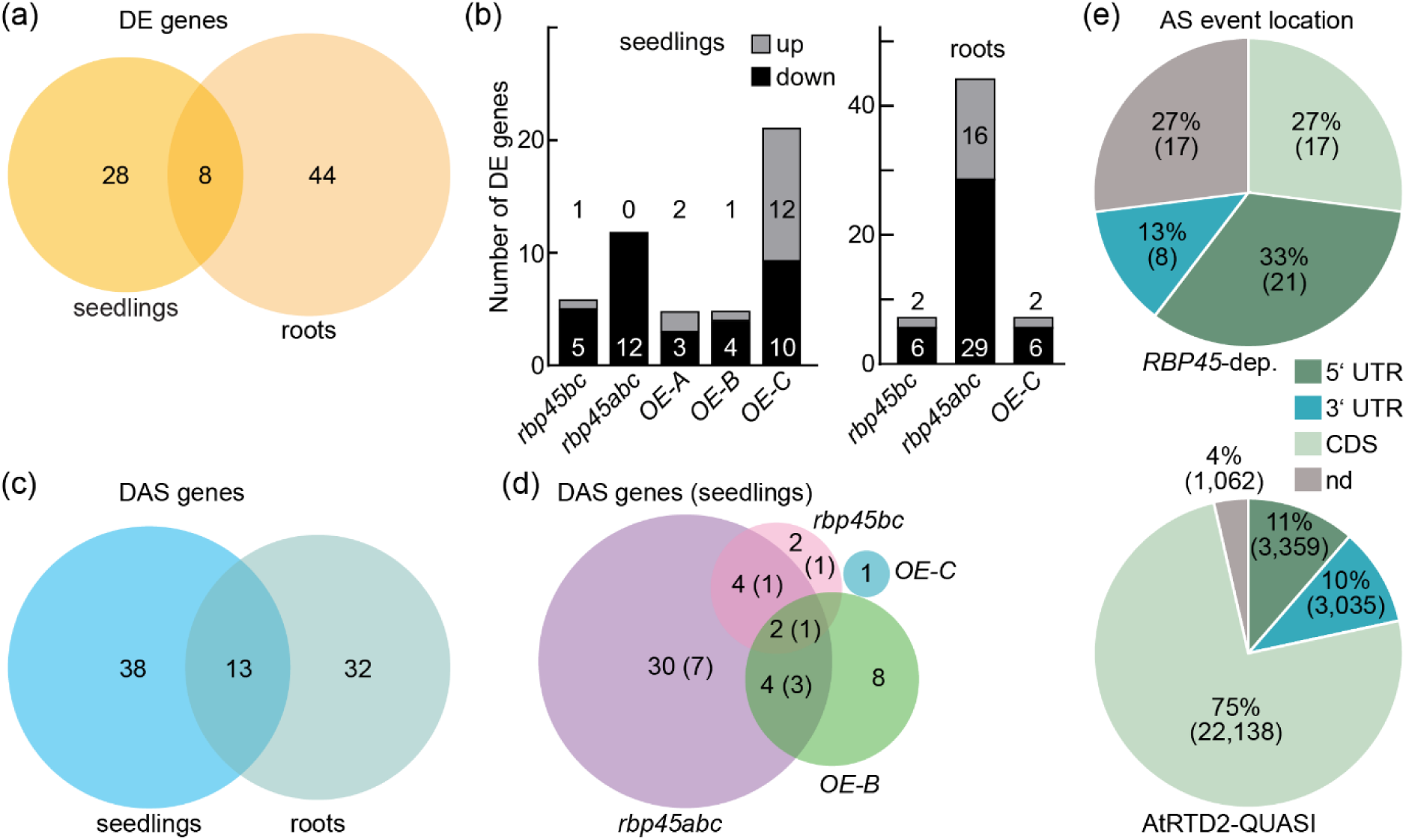
RBP45 misexpression affects expression and AS of few genes in *A. thaliana*. (a, b) Differential gene expression in *rbp45* mutants compared to WT for 10-day-old whole seedlings or roots from 7-day-old plants. Venn diagram (a) shows numbers of combined significant genes, while bar plot (b) provides line-wise comparisons and direction of changes. (c, d) Genes showing differential AS for samples as defined in (a, b). Venn diagram of DAS genes cumulated for seedling and root samples (c) or for individual mutant comparisons in seedlings (d) are displayed. Numbers correspond to DAS genes, with number in parenthesis in (d) indicating the overlap with the root data. (e) Positioning of DAS events (RBP45-dep., top) compared to all events (AtRTD2-QUASI, bottom) in proportions and total numbers (in parentheses). Based on their location relative to translational start and stop sites, events were assigned to the 5’ UTR, CDS, or 3’ UTR; “nd” refers to events that were overlapping or could not be located, as further described in Figure S9c and the method section.

The separate DE analyses for seedling and root samples indicated that only few genes are regulated in an *RBP45*-dependent manner. We next extended our analysis to the line-wise comparison of tissue-specific expression patterns, which might differ between the mutants and the WT and could have been missed in the previous comparisons within one tissue type due to small changes. As we used in our study different number of replicates for the seedling and root samples, we first performed pair-wise comparisons for the WT and three types of mutants (Figure S8a-d). The comparison of the replicates and genotypes gave consistent results, with an overlap of 4,277 (68.4%) DE genes that show differential expression in seedlings and roots of both WT and the two knockout mutant backgrounds *rbp45bc* and *rbp45abc* (Figure S8e). Additionally including the *OE-C* line reduced the number of common DE genes only slightly (Figure S8f; 4,100 genes, 63.2%), representing a robust signature of tissue-specific gene expression that showed in a GO term analysis among others an enrichment of photosynthesis-related genes (Supplemental Data Set 3), as expected in the comparison of phototrophic seedlings and heterotrophic roots. The DE genes not shared between the WT and mutants might point at processes that are disturbed upon RBP45 misexpression and therefore could contribute to the altered root growth. GO analysis for the corresponding gene sets showed no significant enrichment. Further research will be needed to validate these possible differences in tissue-specific gene expression and to examine whether those are linked to the altered root growth upon RBP45 misexpression.

Next, we determined the numbers of differentially alternatively spliced (DAS) genes by comparing the mutant lines to WT, resulting in the seedling and root datasets, respectively, in 51 and 45 affected genes (Figure 8c and Supplemental Data Set 2). Considering the AS changes upon RBP45 overexpression, *RBP45B* resulted in most changes among the three corresponding mutants with 14 DAS genes in whole seedlings (Figure 8d). No DAS genes were detected for the *RBP45A* overexpression seedlings, whereas the *RB45A* gene was the only DAS target in the *OE-C* line, further supporting the cross-regulatory mechanism (Figure 8d). For seedlings of the knockout mutants *rbp45bc* and *rbp45abc*, respectively, 8 and 40 DAS genes were detected, with 6 DAS genes shared between the two mutants (Figure 8d). A similar extent of AS changes was found in the root dataset, with 13 and 39 DAS genes, respectively, in the double and triple mutant, and 7 overlapping genes (Figure S9a). Upon *RBP45C* overexpression, two DAS genes were found in the roots, of which one each overlapped with the dataset for the *rbp45bc* and the *rbp45abc* mutant. To further characterize the alternative splicing changes, we assigned differential transcript usage (DTU) transcripts to the identified DAS genes. In cases where expression levels of two transcripts from the same gene were significantly and reciprocally altered in at least one type of misexpression line compared to the WT, AS event types and their genomic locations were determined (Supplemental Data Set 2). This annotation was performed using the *A. thaliana* TAIR 10 genome and AtRTD2-QUASI reference transcriptome. Considering the AS event positions relative to the open reading frame, approximately half of the significantly altered events were located in the 5′ and 3′ UTR (Figure 8e and S9). This enrichment compared to the reference data set suggests that RBP45-mediated AS to a major extent affects non-coding regions.

To validate the transcriptome-wide findings, RT-PCR analyses were performed for seven candidate DAS events (Figure 9 and S10). Out of those, three events were significantly changed in both RNA-seq datasets (*AVT6*, *HYH,* and *VAB2*), three (*ABC1K8*, *DEP1,* and *ALAD1*) were unique to whole seedlings and one (*XBAT35*) was only detected in the root dataset. According to the RNA-seq data, only *DEP1* and *AVT6* exhibited significant reciprocal splicing changes in comparison of the two knockout lines and the *OE-B* lines. In our validation experiments, all seven AS events were tested across all genotypes, and the significant changes from the RNA-seq were independently confirmed (Figures 9 and S10). In case of *rbp45bc* and *rbp45abc,* all candidate events were significantly and consistently altered both in the two mutants and the two tissue types. With respect to the quantitative change, the effect was slightly but consistently more pronounced in the triple mutant, suggesting an additive effect of RBP45 paralog loss. Interestingly, an in general opposite AS shift was seen for *OE-B*, while overexpression of the other two RBP45 paralogs did not have such an effect (Figure 9 and S10). This further supports a prominent function of *RBP45B* in AS regulation among the RBP45 family, and is consistent with its central role in auto- and cross-regulation.

**Figure 9:**
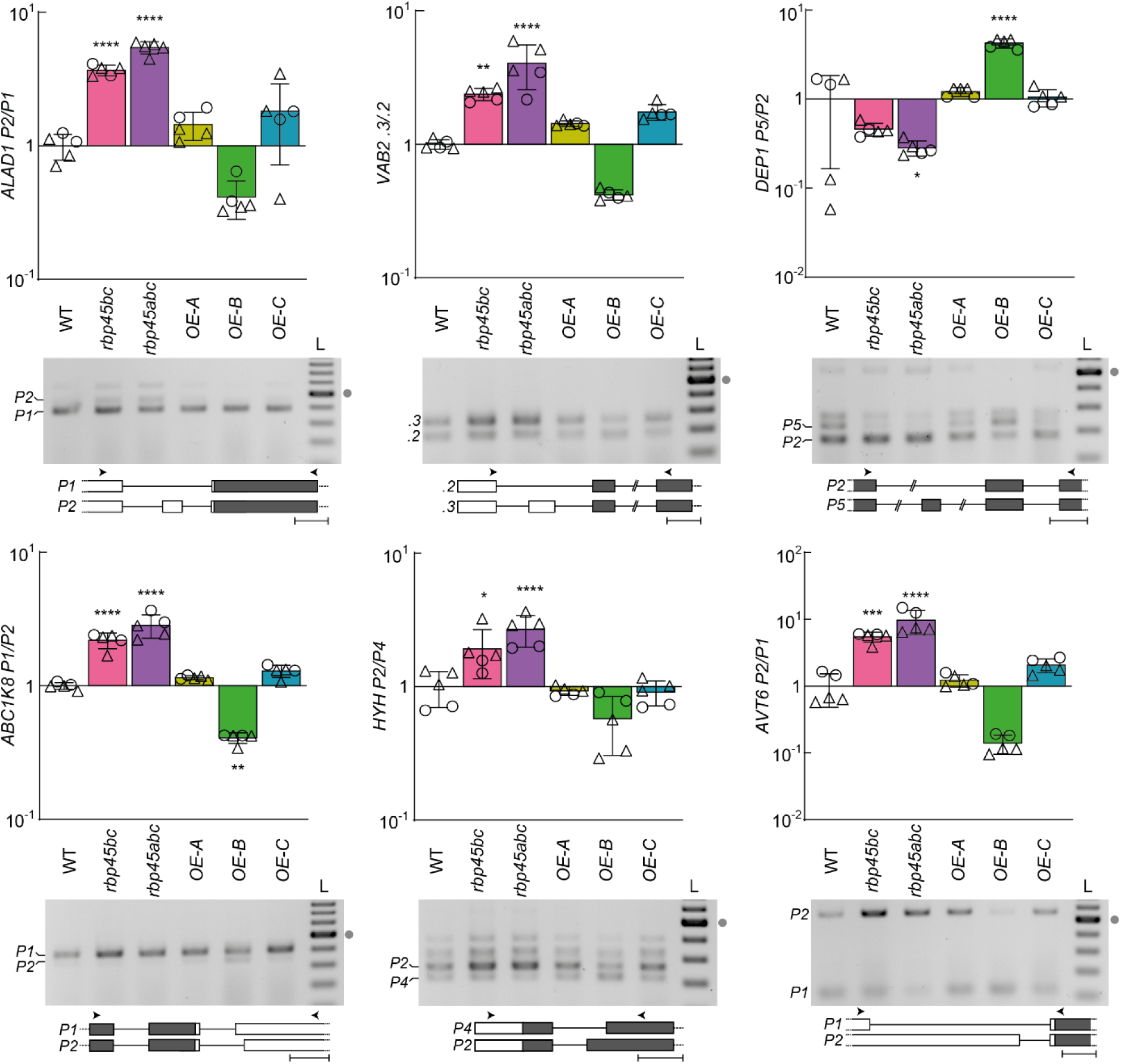
Reciprocal AS shifts upon *RBP45B* knockout and overexpression confirm its function in splicing regulation. AS ratios were analysed from 10-day-old *A. thaliana* WT, *rbp45bc, rbp45abc, OE-A#15, OE-B#1, and OE-C#8* for *ALAD1* (*AT1G69740*), *VAB2* (*AT4G38510*), *DEP1* (*AT5G53850*), *ABC1K8* (*AT5G64940*), *HYH* (*AT3G17609*), and *AVT6* (*AT3G30390*). For each AS event, bar chart based on Bioanalyzer quantification (top) and representative gel picture (middle) of RT-PCR co-amplification products, and the corresponding gene models (bottom) are displayed. Mean value (bars), standard deviation (error bars), and individual data points (circles: based on samples used for RNA sequencing; triangles: additional replicates) are depicted each; mean AS ratio of WT was set to 1. Asterisks indicate significant change compared to WT (one-way ANOVA followed by Dunnett’s multiple comparisons test, *p < 0.05, **p < 0.01, ***p < 0.001, ****p < 0.0001). Size ladder (L) for gels consisted of DNAs in 100 bp increments with the strongest band corresponding to 500 bp (marked with gray dot). In the gene models, exons, introns, CDS, and UTRs are depicted by boxes, lines, black, and white shading, respectively; double dashes indicate cropped regions; arrowheads show primer binding sites and scale bar is individually adjusted to 100 bp for each model.

## Discussion

### Crosstalk of RBP45 homologs via AS regulation

Here, we characterized the role of the strucRNA *45ABC* in AS regulation of *RBP45A*, *RBP45B*, and *RBP45C* from *A. thaliana*. Our results revealed negative auto- and cross-regulation of these three genes via AS, balancing the ratio of protein-coding *cd* variants to non-productive *nc* variants that are targeted by NMD (Sack *et al*., 2025). This regulatory mechanism is common among RBP genes and enables rapid and dynamic fine-tuning of protein levels (Müller-McNicoll *et al*., 2019; Reddy *et al*., 2013; Wachter *et al*., 2012). In line with autoregulation, constitutive overexpression of any RBP45 member led to reduced levels of the *cd* isoform for the corresponding endogenous gene (Figure 3). With respect to cross-regulation, *RBP45B* seems to play a dominant role among the three homologs. Its overexpression resulted in an increase of the *nc* variants for all three RBP45 genes, whereas in lines overexpressing *RBP45A* or *RBP45C* both the *cd* and *nc* isoform levels from all three endogenous RBP45 genes were diminished (Figure 3b, c). Accordingly, formation of the non-coding variants from these genes is to a major extent caused by *RBP45B*, which in turn is suppressed in the *RBP45A* and *RBP45C* overexpression lines. This regulatory hierarchy is further supported by the analysis of the knockout mutants (Figure 4). Knocking out only *RBP45B* reduced *nc* isoforms for all three genes, whereas total levels of *RBP45A* and *RBP45C* transcripts were elevated. In the *rbp45bc* double mutant, however, *RBP45B nc* levels were restored to a WT-like level. In parallel, the AS pattern of *RBP45A* was altered in the double mutant, with an increase of *cd* and decrease of *nc* transcripts, pointing towards a release of negative cross-regulation by *RBP45C*. As a consequence, *RBP45A* was de-repressed in the double mutant, and triggered enhanced production of *nc* transcripts from the *RBP45B* gene (Figure S11). Moreover, the *rbp45abc* triple mutant had strongly diminished amounts of *nc* isoforms for all three genes, suggesting that this regulatory circuit is probably confined to the three RBP45 genes.

Similar auto- and cross-regulatory circuits have been identified in plants before, including studies of the hnRNP genes *GRP7* and *GRP8* (Schöning *et al*., 2007; Schöning *et al*., 2008) and the three *PTB* genes (Rühl *et al*., 2012; Stauffer *et al*., 2010) from *A. thaliana*. AS-based cross-regulation was observed for *GRP7*/*GRP8* and *PTB1*/*PTB2*, whereas the more distantly related *PTB3* had no effect on the splicing outcome of its two paralogs. What is then the molecular mechanism underlying the hierarchy of the more complex regulatory circuit shown in this study for the RBP45 genes? And why does it involve a conserved structured mRNA motif as opposed to a simpler pre-mRNA architecture containing single or several binding sites for the corresponding RBP as has been demonstrated in other instances?

Evolutionary analyses indicate that *RBP45B* is the most ancient one of the three paralogs, while *RBP45C* and *RBP45A* emerged more recently and are more closely related (Lorković *et al*., 2000; Park *et al*., 2006). This relationship is also reflected by the different domain organization of the proteins. All three RBP45 proteins contain three RRMs; while RBP45A and RBP45C have in addition a glutamine-rich N-terminus, RBP45B is enriched in glutamine residues near the C-terminus and exhibits a proline-rich N-terminus. These differences might be responsible for or contribute to the distinct roles of the three RBP45 genes in AS regulation. Moreover, varying total and spatial expression patterns might play a role, and we observed in WT plants that the transcript levels for *RBP45B* were highest among the three homologs (Supplemental Data Set 2). A major role of *RBP45B* in AS regulation is not only supported by its function in the auto- and cross-regulatory circuit, but also our transcriptome-wide analyses. *RBP45B* overexpression caused the highest number of significant AS changes (Figure 8) as well as reciprocal AS shifts compared to the double and triple knockout mutants (Figure 9). Given that even for the *RBP45B* overexpression line and the triple mutant *rbp45abc* relatively few global AS changes were detected, redundancy with other RBPs and/or additional functions of these RBP45 genes besides AS regulation can be assumed. Moreover, our observation of specificity among the RBP45 genes indicates unequal genetic redundance, which has been previously discussed to be a more common evolutionary process of functional gene diversification in *A. thaliana* (Briggs *et al*., 2006).

### Critical and interconnected roles of *45ABC* sequence and structure in balancing RBP45 expression

The exclusive presence of the strucRNA *45ABC* in the three RBP45 genes and the associated cross-regulatory network suggested that their expression is coordinated in an RNA structure-dependent manner. RNA folds can affect RBP interactions by various means, e.g. through exposing or occluding binding sites, while RBP binding in turn can remodel the RNA structure. Recent transcriptome-wide structure-profiling studies in plants further highlighted the critical interplay between RNA sequence and structural features. For example, Gosai *et al*. (2015) observed that RBP binding sites in nuclear mRNAs are generally less structured, and Liu *et al*. (2021) reported that the 5’ splice site and branch point need to be single stranded for efficient splicing. In plants, few examples exist where the impact of RNA structure and its dynamical behaviour on AS or other steps of gene expression has been functionally characterized. One such case is the TPP riboswitch mechanism that functions via occlusion of the alternative 5’ splice site by the interaction with an aptamer region in the absence of ligand binding (Wachter *et al*., 2007). AS regulation via changes in splice site availability was also reported for the heat shock factor *HsfA2* gene from tomato, involving temperature-dependent structural changes at a 3’ splice site (Broft *et al*., 2022). An alternative mechanism has been proposed in AS control of the *TFIIIA* gene via the strucRNA *P5SM* (Hammond *et al*., 2009), where L5 binding may prevent binding of a splicing regulatory protein independent of an RNA structural change.

Using a mutational approach, we showed in this work that opening the structure of *45ABC* resulted in increased usage of the alternative splice sites (Figure 5). The most pronounced shift towards the *nc* isoform including the CE was observed upon disruption of stem I. Compensatory mutations that restored base-pairing also rescued the splicing pattern to a WT-like AS ratio, underscoring the structural requirement for proper splicing regulation by this element. The important role of stem I in AS regulation combined with the localization of the alternative 5′ splice site and the respective U1 snRNP-binding motif (Sheth *et al*., 2006) within stem II suggest that the assembly of the two hairpins is required for proper splice site selection. Sequence and structure conservation of *45ABC* may therefore be explained by its several roles; as a binding site for RBP45 and U1 components, its structural constraints, and potential crosstalk between the elements. Notably, all three RBP45 proteins have been shown to interact with the U1 snRNP component U1-C (Huang *et al*., 2022). Their human ortholog TIA-1 is described to bind downstream of the 5’ splice site and to recruit U1 snRNP through a direct interaction with U1-C, thereby influencing splice site selection (Förch *et al*., 2002). In the case of the RBP45 proteins, their binding to stem I of *45ABC* might facilitate U1 recruitment to the alternative 5’ splice site embedded in stem II and thereby promote its usage. Accordingly, this strucRNA may act as a sensor for RBP45 protein levels that allows balancing RBP45 expression in an AS-dependent manner (Figure 10). Closely related proteins, namely RBP45D and RBP47 (Chang *et al*., 2022), could potentially also bind *45ABC* and affect the splicing outcome. RBP45D has been reported to be involved in alternative splice site selection via interaction with U1 components (Chang *et al*., 2022), whereas the RBP47 proteins have been described as regulators of stress granule dynamics and therefore may primarily act in other aspects of RNA metabolism (Kosmacz *et al*., 2019). Functional redundancy between RBP45 and possibly also RBP47 proteins in U1 recruitment to 5’ splice sites could also explain why even in the triple mutant *rbp45abc* only relatively few AS changes were detected in our transcriptome-wide analysis. Besides being part of the spliceosome, U1 snRNP also performs splicing-independent functions. In a process termed telescripting, U1 binding blocks recognition of cryptic polyadenylation signals and thereby prevents premature cleavage and polyadenylation (Berg *et al*., 2012; Kaida *et al*., 2010). This additional U1 snRNP function has recently also be demonstrated for *A. thaliana* (Mangilet *et al*., 2024) and future experiments should address a possible involvement of RBP45 proteins.

**Figure 10:**
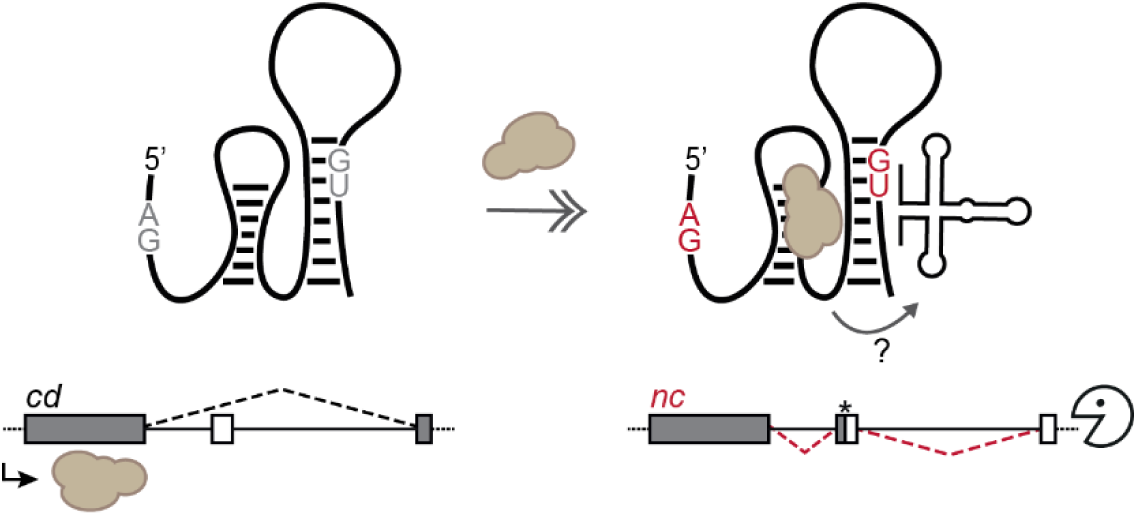
Model of *45ABC*-mediated AS regulation. The strucRNA *45ABC* consists of two stem loops, with the second one encompassing the alternative 5’ splice site used for CE inclusion. When RBP45 protein (brown cloud) is not bound to stem I (left), the CE is removed resulting in the *cd* variant that is translated into RBP45 protein. RBP45 binding to stem I may facilitate U1 snRNP recruitment to the alternative 5‘ splice site in stem II (right), thereby promoting its usage. The CE-containing *nc* variant contains a premature termination codon (asterisk), triggering NMD turnover as part of the negative feedback regulatory loop. Gray and white boxes in the transcript models correspond to coding and non-coding regions.

We also examined if the conserved purine-rich motif (RGRG) within stem I might serve as a binding site for the three RBP45 proteins and is important for the regulation. In line with this hypothesis, transient reporter assays revealed that mutating the five consecutive guanines to adenines caused an almost complete loss of the *nc* isoform under control conditions (Figure 6). Upon co-expression of any *RBP45* CDS, a splicing response was still detectable, but it was markedly attenuated compared to the WT construct. This residual responsiveness may be explained by RBP45 binding to the introduced U-stretch, consistent with previous findings that recombinantly expressed RBP45 from *N. plumbaginifolia* (Lorković *et al*., 2000) and RBP45D from *A. thaliana* (Huang *et al*., 2022) interacted with U-rich RNAs *in vitro*. Alternatively, or additionally, it might reflect non-specific interactions caused by high levels of RBP45 expression in the transient expression system. With regard to the proposed binding preference of the RBP45 proteins, the U-rich loop of stem I (Sack *et al*., 2025) may also contribute to RBP45 binding. Furthermore, it is well known that RBPs differ not only in their sequence preferences but also with respect to the structural requirements at the binding site and its context (Dominguez *et al*., 2018; Foley *et al*., 2017; Gosai *et al*., 2015). Accordingly, conservation of the stem I structure of *45ABC* might be a consequence of the RBP45 proteins’ binding preferences. Taken together, both sequence and structural features of *45ABC* contribute to this RBP45-responsive regulatory mechanism, providing also an explanation for the relatively low level of covariation of this motif (Sack *et al*., 2025).

Our findings raise the question whether related mechanisms of strucRNA-dependent AS are more common. Interestingly, Sack *et al*. (2025) also identified conserved strucRNAs within the pre-mRNAs of *GRP7*, *GRP8*, and *PTB2.* The *GRP7&8* strucRNA consists of a hairpin including an alternative 5’ splice site in the predicted stem (Sack *et al*., 2025). Usage of this splice site leads to NMD-sensitive isoforms as part of negative feedback regulation (Schöning *et al*., 2008), and both GRPs have been shown to bind downstream of this motif (Leder *et al*., 2014; Staiger *et al*., 2003). Accordingly, the corresponding strucRNA may suppress usage of this splice site, while GRP binding could promote it (Sack *et al*., 2025). Similarly, the *PTB2* motif is a single hairpin that includes the 5’ splice site and a polypyrimidine stretch that are used for AS-mediated negative auto- and cross-regulation of *PTB2* expression (Burgardt *et al*., 2024; Sack *et al*., 2025; Stauffer *et al*., 2010). So far experimental evidence for the functional relevance of these novel strucRNAs is lacking. However, their discovery suggests that such RNA folds might be more common in AS regulation. Reasons for their sequence and structural conservation can include the presence of binding sites for splicing factors and regulators, specific requirements for structuredness, and regulatory interactions between individual elements as proposed for *45ABC*. Shorter motifs such as single hairpins are more difficult to detect by bioinformatics approaches searching for conserved structural elements and thus might be more common than reflected by our current knowledge.

### Physiological roles and functions of RBP45 proteins

Our transcriptome-wide analyses of *rbp45* knockout and overexpression lines revealed only limited changes in global gene expression and AS. Interestingly, most observed AS changes occurred in UTRs (Figure 8 and S9). This enrichment could point at additional functions of RBP45 association with its target RNAs that occur downstream of AS control, such as the established role of UTR-binding proteins in processes such as mRNA translation, localization, and stability (Hardy and Balcerowicz, 2024). Consistent with such an idea, RBP45 proteins localize to both the nucleus and cytoplasm (Huang *et al*., 2022; Lorković *et al*., 2000) and interact with mRNA decay components such as UPF1 (Chicois *et al*., 2018; Sulkowska *et al*., 2020). Protein-protein interaction studies further support a role of RBP45 proteins in regulating the mRNA fate, e.g. RBP45B was shown to interact with the cap-binding protein CBP20 and the poly(A)-binding protein PAB8 (Muthuramalingam *et al*., 2016). Furthermore, RBP45A, RBP45B, and RBP45C were reported to be associated with the UTRs of the *GRP7* mRNA (Reichel *et al*., 2024). These interactions and the concomitant regulatory potential might be particularly important under stress. Accordingly, expression of *RBP45A* and *RBP45B* has been reported to be induced upon ozone exposure (Peal *et al*., 2011).

Critical functions of the RBP45 genes in plant development are supported by the mutants’ phenotypes. With an increasing number of knocked out RBP45 genes, we observed a progressively reduced primary root length (Figure 7a). This observation also suggests functional redundancy among the three paralogs. Interestingly, overexpression of *RBP45C* similarly disturbed root growth. Inspecting the transcript levels in this overexpression line from the RNA-seq data revealed a massive overaccumulation of *RBP45C* specifically in the root, pointing at tissue-specific regulation (Supplemental Data Set 2). This overaccumulation may have caused suppression of the other two RBP45 paralogs and therefore resulted in a similar phenotype as for the knockout lines. Further research is needed to define the responsible target genes or processes. Interestingly, a previous study by Foley *et al*. (2017) identified an interaction of RBP45 proteins with a TG-rich motif in the 3’ UTRs of root hair-specific transcripts, potentially linking RBP45 function to root cell fate decisions. Beyond changes in root length, *rbp45* mutants showed altered flowering time. Overexpression of *RBP45A* caused a mild delay, while *RBP45B* and *RBP45C* overexpression led to earlier and later flowering, respectively (Figure 7b). Delayed flowering upon loss of *RBP45B* is consistent with previous observations (Muthuramalingam *et al*., 2016). Interestingly, this phenotype was absent in higher-order mutants, suggesting a potential antagonism or compensatory functions among *RBP45* homologs in this process.

In conclusion, our work has identified novel physiological functions of the RBP45 genes in plant development. Furthermore, we have characterized an intricate auto- and cross-regulatory circuit that balances the expression of the three RBP45 genes via the strucRNA *45ABC*. The conserved sequence and structural features of this element point towards an AS regulatory mechanism that involves RBP45-mediated recruitment of U1 snRNP to an alternative splice site, resulting in negative feedback control of RBP45 expression.

## Materials and methods

### Plant cultivation and transformation

*Arabidopsis thaliana* ecotype Columbia-0 plants were either grown on soil or in sterile culture on agar plates. Therefore, seeds were surface-sterilized using 3.75% NaClO and 0.01% Triton X-100 and stratified at 4 °C for 2 - 4 days. If not specified otherwise, they were germinated on ½ Murashige and Skoog medium including vitamins (Duchefa) and containing 2% sucrose and 0.8% phyto agar. Segregating mutant lines were grown on plates containing additionally 25 µg/mL kanamycin. For experiments in which root lengths were measured, 1.2% phyto agar was used and plates were placed not in horizontal but vertical orientation. The standard settings of the climate chambers were 16 h white light (∼100 µE)/22 °C and 8 h darkness/20 °C at 60% relative humidity. *Nicotiana benthamiana* was grown on soil in a climate chamber (16 h white light (∼120 µE)/24 °C, 8 h dark/22 °C at 60% relative humidity).

Stable transformation of *A. thaliana* was achieved by the floral dip method described by Clough and Bent (1998). Selection of primary transformants was done on half-strength Murashige and Skoog medium (including vitamins) containing 0.8% plant agar, 25 µg/mL kanamycin, and 200 µg/mL cefotaxime. Resistant plants were transferred to soil and upon further growth subjected to PCR-based genotyping. Selection of CRISPR lines was carried out via the fluorescence-accumulating seed technology (FAST; Shimada *et al*., 2010) under a Leica M205FCA fluorescence stereomicroscope (excitation: 470 nm, emission: 585 nm) and positive seeds directly germinated on soil. Cas9-free plants in later generations were confirmed by the absence of fluorescence.

*Nicotiana benthamiana* leaves were transformed via Agrobacteria-mediated leaf infiltration as described by Wachter *et al*. (2007) with a normalization construct based on mOrange2 (Shaner *et al*., 2008). After two days, leaf material was harvested two and three days after infiltration, respectively, for analysis on RNA and protein level.

### Genotyping of plant mutants

The genetic status of the generated mutants was confirmed by isolating genomic DNA followed by genotyping PCR. To 100 mg ground plant tissue, 500 µL extraction buffer (200 mM Tris-HCl pH 9, 400 mM LiCl, 25 mM EDTA, 1% SDS) was added and vortexed thoroughly at room temperature. After 5 min centrifugation at 15 000 *g*, the supernatant was mixed with the same volume of isopropanol, incubated for 5 min, and then centrifuged for 10 min at 15;000 *g* to precipitate the gDNA. The pellet was washed with 70% Ethanol, dried at 37 °C, and resuspended in ½ TE buffer (5 mM Tris-HCl pH 8, 0.5 mM EDTA). Genotyping PCR was performed with homemade Taq polymerase according to standard procedures with primers listed in Supplemental Data Set 4. For CRISPR lines, a proof-reading polymerase was used for PCR amplification, followed by purification of PCR fragments (GeneJET PCR Purification Kit, Thermo Fisher Scientific) and Sanger sequencing of mutated sites.

### Plant phenotyping

Seeds were sorted using Boxeed 2.1 (Labdeers) with the following criteria: pixel count range 200 – 2000, SSE 0 – 95, and LS ratio 0 – 25. Measurements and pictures of matching seeds were taken, and size distribution was analyzed.

For root length measurements, seeds were also stratified for 96 h in darkness at 4 °C before plating them on square ½ MS plates singly in one row. The plates were placed vertically into racks and kept under beforementioned conditions for 7 days. The plates were scanned, and primary root length was measured using ImageJ software (Schindelin et al. 2012).

For the determination of flowering time, seeds were stratified for 96 h in darkness at 4 °C and then cultivated on soil under the conditions mentioned above. Flowering time was defined as the time span at which the inflorescence reached a length of 1 cm.

### Cloning procedures

The splicing reporters for *RBP45A* (*AT5G54900*), *RBP45B* (*AT1G11650*), and *RBP45C* (*AT4G27000*) were constructed by PCR amplification of the corresponding sequences from *A. thaliana* genomic DNA using the oligonucleotides indicated in Supplemental Data Set 4 and sub-cloning into pGEM-T (Promega, www.promega.de). All final reporter constructs were cloned in the pBinAR vector (Höfgen and Willmitzer, 1990) using the previously published *TFIIIA* reporter (Hammond *et al*., 2009) by replacing the target gene region via *Kpn*I/*Xba*I restriction digest. Motif mutations were introduced via PCR mutagenesis in pGEM-T using oligonucleotides as indicated in Supplemental Data Set 4. The CDS constructs used for leaf infiltration assays and generating the stable constitutive overexpression lines were also cloned in the pBinAR vector. Upon PCR amplification from cDNA, the fragments were first subcloned into pGEM-T and subsequently ligated into pBinAR via *Kpn*I/*Xba*I. All final constructs were confirmed by sequencing of the inserts. The CRISPR-Cas9 constructs for generation of *rbp45* knockout plants were created using the GoldenGate cloning system reported by Stuttmann *et al*. (2020). The sgRNAs indicated in Supplemental Data Set 4 were designed using CHOPCHOP (Labun *et al*., 2019) and cloned into the pDGE652 or pDGE347 vector under control of the *A. thaliana U6-26* promoter fragment as described by Stuttmann et al. (2020). The vector additionally features an intron-containing *Cas9* under control of the *RPS5A* promoter.

### RNA isolation & analysis of splicing patterns

RNA was isolated from ∼100 mg ground plant tissue using EURx Universal RNA Purification Kit (Roboklon) with an on-column DNase treatment (RNase free DNaseI, NEB) as specified in the manufacturer’s protocol. The RNA concentration was determined using a spectrophotometer (DeNovix DS11 FX+). Reverse transcription was carried out with SuperScript II (Invitrogen) using a dT_20_ primer with a 5 min incubation step at 65 °C prior to adding the reaction buffer and enzyme. The cDNA obtained was subjected to analysis of splicing patterns via co-amplification PCR or quantitative PCR (qPCR).

Co-amplification PCR was performed with primers (Supplemental Data Set 4) encompassing the alternative spliced regions and a homemade Taq polymerase. The resulting DNA products were separated on horizontal agarose gels. Either a 1 kb plus (New England Biolabs) or a 100 bp plus GeneRuler ladder (Thermo Scientific) was used to estimate the molecular weight. The bands were stained after electrophoresis in ethidium bromide solution at 0.5 µg/mL. The bands were visualized with UV light and documented with the Quantum-CX5 Edge (Vilber). Quantification of co-amplification PCR products was performed with a Bioanalyzer 2100 (Agilent, www.agilent.com) using the DNA1000 chip according to manufacturer’s protocol.

qPCR was performed using the Biorad CFX384 Real-Time PCR system (Biorad, www.biorad.com) and the MESA-BLUE 2x qPCR MasterMix Plus for SYBR® Assay (Eurogentec, www.eurogentec.com). Primers are listed in Supplemental Data Set 4 and primer efficiencies were determined with serial dilutions of template. All reactions were done in triplicates, and a melting curve analysis was included. Analysis of the data was done with the use of the relative standard curve method. Transcript level of *PP2A* (*AT1G13320*) served as reference.

### Whole transcriptome sequencing

150 ng total RNA isolated as described before and derived from two biological replicates of 10-day old *A. thaliana* WT, *rbp45abc, rbp45bc,* and *RBP45A*/*RBP45B/RBP45C* overexpression seedlings was utilized. For root-specific analysis, RNA was used from clipped roots of 7-day-old seedlings with three biological replicates each for WT, *rbp45bc*, *rbp45abc*, and the *RBP45C* overexpression line. The concentration and quality of the RNA was assessed based on the RNA integrity number (RIN) measured with an Agilent Bioanalyzer 2100 using the RNA6000 Nano protocol. Eurofins Genomics performed poly-A-selection and generation and sequencing of strand-specific cDNA libraries on a NovaSeq 6000 S4 system, generating a minimum of 25 million read pairs (2 x 150 nt) per sample. BBDuk (Bushnell B., www.sourceforge.net/projects/bbmap) was used for trimming the reads, which were then mapped to the reference transcriptome AtRTD2-QUASI (Zhang *et al*., 2017) using Salmon (Patro *et al*., 2017). Data was analyzed using the 3D RNA-seq App (Guo *et al*., 2021). Further details are given in Supplemental Methods.

To determine relative positions of AS events within transcripts, first the longest possible ORF from each transcript in AtRTD2-QUASI was determined. For each gene, the transcript with the longest ORF was set as reference to define coordinates of translation start and end sites. Based on these coordinates, AS events were localized in all transcript isoforms per gene. Only the first event from the 5’ end was considered. The same events in multiple isoforms were counted only once. Further details are provided in Supplemental Methods.

Venn diagram displays were created with DeepVenn (Hulsen, 2022). Gene ontology term analysis was performed using ShinyGO 0.85 (Ge *et al*., 2020) with as FDR cutoff of 1 x 10^-5^ using the KEGG pathway database (Kanehisa *et al*., 2021).

### Protein extraction and fluorescence assay

Total protein was isolated from ∼100 mg ground tissue of infiltrated *N. benthamiana* leaves. After adding 300 µL buffer (50 mM Tris pH 7.5, 150 mM NaCl, 0.1% Tween, 0.1% ß-mercaptoethanol), the samples were vortexed and centrifuged at 4 °C for 15 min at 15 000 *g*. Supernatant was transferred to a new tube and then 100 µL of each sample pipetted into a 96 well-plate (Greiner, black, flat bottom) for fluorescence measurement. The fluorescence was detected using the TECAN infinite M1000, with excitation at 478 - 492 nm and emission at 515 - 525 nm for EGFP measurement. For fluorescence measurement of the reference mOrange2, the settings 525 - 530 nm for excitation and 590 - 610 nm for emission were used. Measurements were taken with flash frequency of 400 Hz and integration time of 20 µs. The measured values were read out in TECAN i-control and transferred to Microsoft Excel (Microsoft Office 2016) for evaluation.

### Statistical analyses

Statistics were carried out with Prism GraphPad 9.4.0 (GraphPad; www.graphpad.com) or, in case of seed surface data, using R Statistical Software (v4.1.2; R Core Team 2021). It was assumed that data follow a normal distribution. Results from statistical analysis are listed in Supplemental Data Set 5.

## Author contributions

Conceptualization: A.W., M.R, Z.W..; Investigation: M.R., R.B., C.E., M.S.; Writing: M.R. and A.W.; Supervision: A.W. and Z.W.; Funding acquisition: A.W. and Z.W.

## Supporting information

Supplemental data set 1

Supplemental data set 2

Supplemental data set 3

Supplemental data set 4

Supplemental data set 5

## Acknowledgements

We are grateful to Christine Wendler and Celine Denrath for technical assistance.

## Data statement

RNA-seq data have been deposited in the European Nucleotide Archive (ENA) repository (https://www.ebi.ac.uk/ena) under accession number PRJEB102517.

