## Supplemental data and methods for "The structured mRNA element *45ABC* mediates auto- and cross-regulation of RBP45 genes via alternative splicing"

**Reinhardt et al.**

**SUPPLEMENTAL DATA**

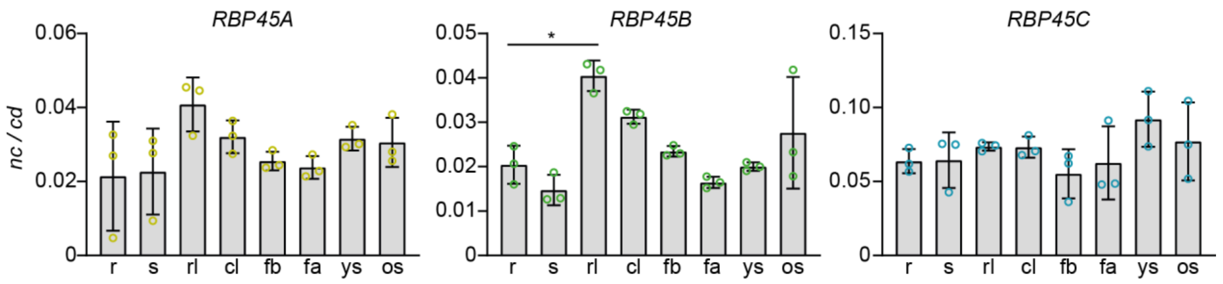


**Figure S1: RBP45 splicing patterns in different tissues.**
Bioanalyzer quantification of AS patterns for *RBP45A*, *RBP45B,* and *RBP45C* in different tissues from *A. thaliana* WT plants. r: seedling roots (7 d), s: total seedlings (12 d), rl: rosette leaves, cl: cauline leaves, fb: flower buds, fa: flower in anthesis, ys: young siliques, and os: old siliques. Circles represent biological replicates (n = 3); displayed are mean values of splicing variant ratios with standard deviation. Asterisk indicates significant difference between tissues; Kruskal-Wallis followed by Dunn's test *p < 0.05.

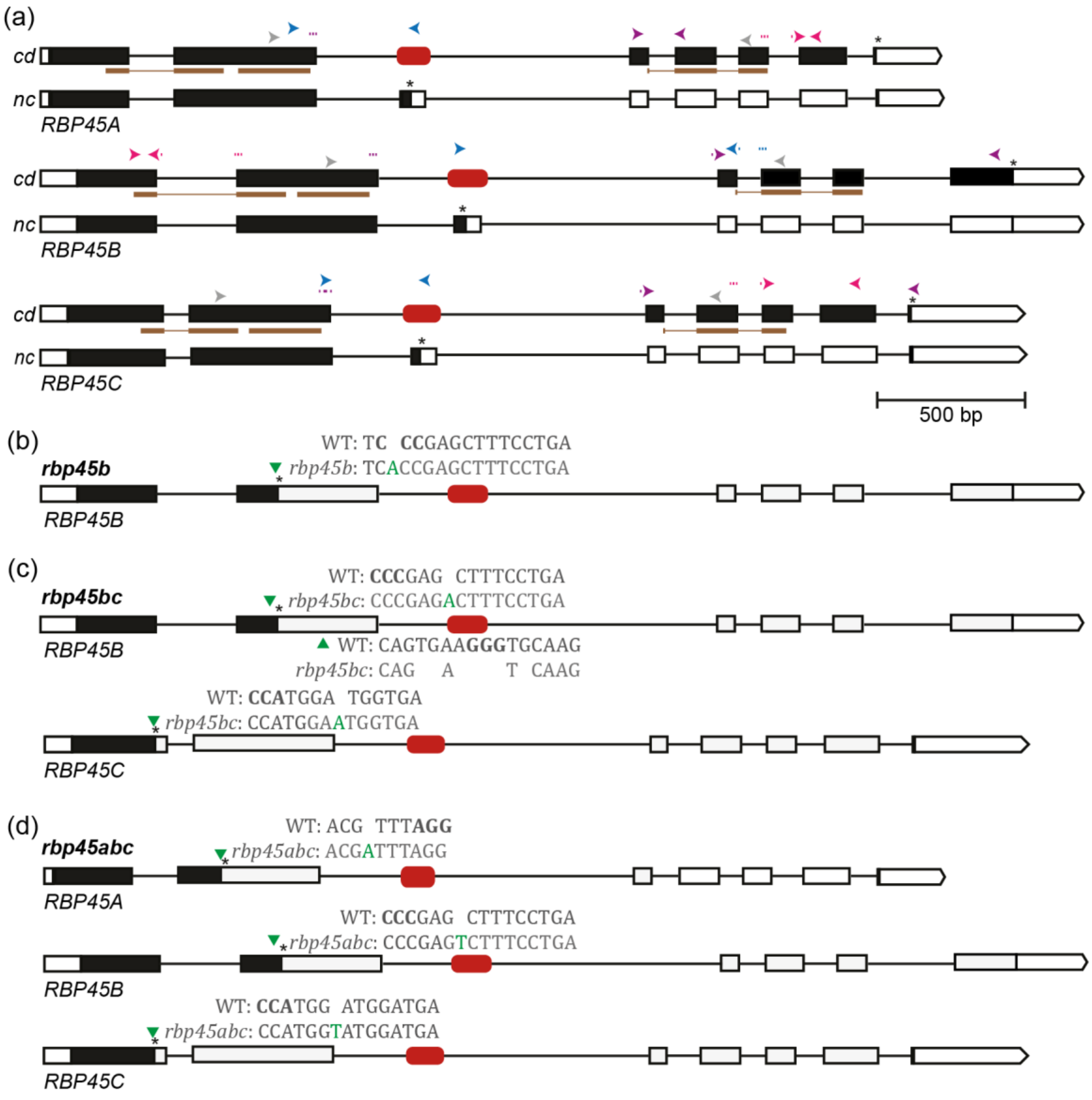


**Figure S2: Models of RBP45 genes and sites of mutations in knockout lines.**
Models of RBP45 genes from *A. thaliana* (a) and CRISPR-induced mutations in *rbp45b* (b), *rbp45bc* (c), and *rbp45abc* (d). Arrowheads show positions and directions of primers used for co-amplification PCR (grey) and qPCR (total levels (pink), non-coding (blue), and coding (purple)); dotted lines depict primers spanning exon-exon borders. Brown lines in (a) show regions coding for RRMs (according to UniProt.org). Exons, introns, CDS and UTRs are depicted by boxes, lines, black and white shading, respectively; *45ABC* motif is indicated by red rounded rectangle. Asterisks mark positions of regular or premature stop codons. Mutation sites are indicated as green triangles, and inserted sequences are shown in green letters; PAM sequence is shown in bold.

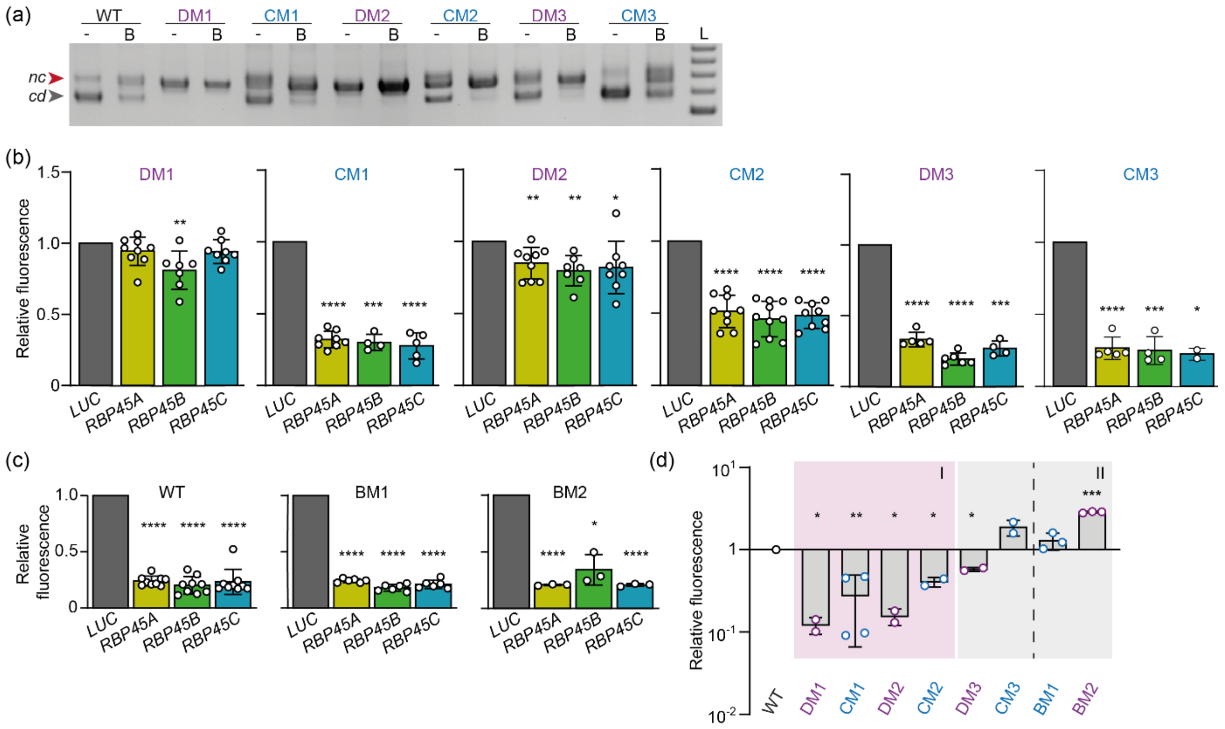


**Figure S3: Mutations within *45ABC* change the splicing outcome of the *RBP45C* reporter upon transient expression in *N. benthamiana*.**

(a) Agarose gel analysis of RT-PCR products of splicing variants from *RBP45C* WT and mutant reporters from *N. benthamiana* leaves co-infiltrated with *LUC* (-) or *RBP45B* (B) constructs. L: size marker, 100 bp increments from 500 – 900 bp.

(b – d) Quantification of reporter output based on GFP fluorescence from transiently transformed *N. benthamiana* leaves co-expressing the indicated splicing reporter with either *LUC* or the CDS constructs of *RBP45A*, *RBP45B*, and *RBP45C* (b, c), or with only LUC construct (d). Mean values and standard deviations are shown; open circles represent individual data points. Data normalization either separately for each reporter to the LUC sample (b, c) or to the mean of WT reporter (d). Asterisks indicate significant differences compared to the LUC control (b, c) or to the WT reporter (d) based on one sample t test (*$p < 0.05$, **$p < 0.01$, ***$p < 0.001$, ****$p < 0.0001$).

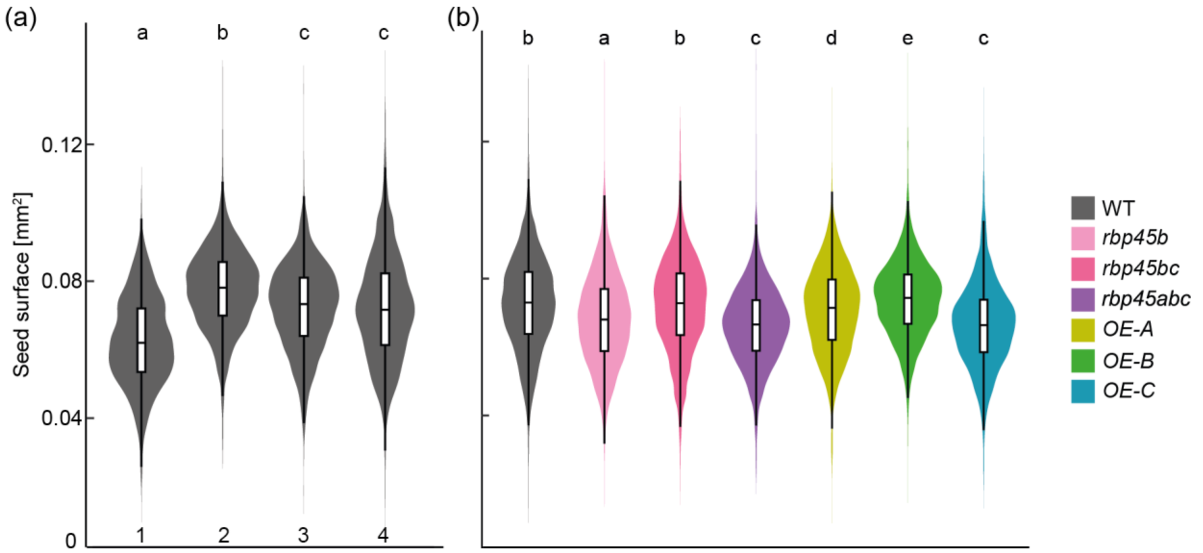


**Figure S4: Variation in seed size is similar in RBP45 misexpression lines and WT.**
Surface area of seeds from *A. thaliana* WT (4 batches from independently grown plants, a) and RBP45 misexpression lines (b). The WT seeds in (b) were a mixture from batches number 2 and 4 from (a). Number of seeds per sample was ≥ 3000 (WT) and ≥ 6000 (*RBP45* misexpression lines). Letters indicate statistically significant differences (ANOVA followed by Tukey test, $p < 0.05$).

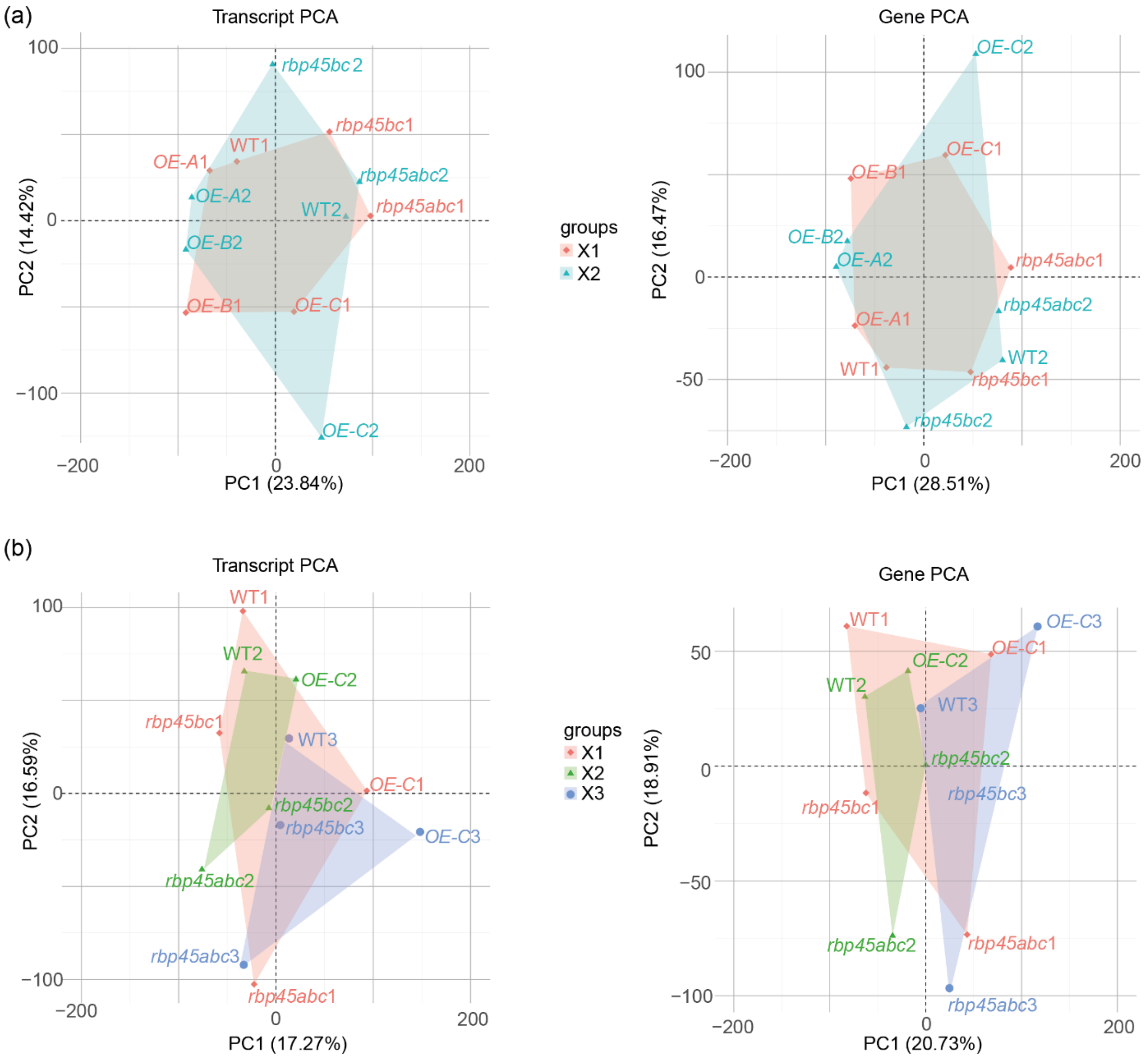


**Figure S5: Principal component analyses of transcriptome datasets.**
Transcript (left) and gene (right) PCA for datasets derived from 10-day-old whole seedlings (a) or roots from 7-day-old seedlings (b) for *A. thaliana* WT, *rbp45bc, rbp45abc*, and constitutive overexpression (OE) lines of *RBP45A, RBP45B*, and *RBP45C*. The suffix in the name refers to the replicate number. Note that the overexpression lines for *RBP45A* and *RBP45B* were only included in the whole seedling dataset in (a). The graphs were created as part of the data analysis in 3D RNA-seq.

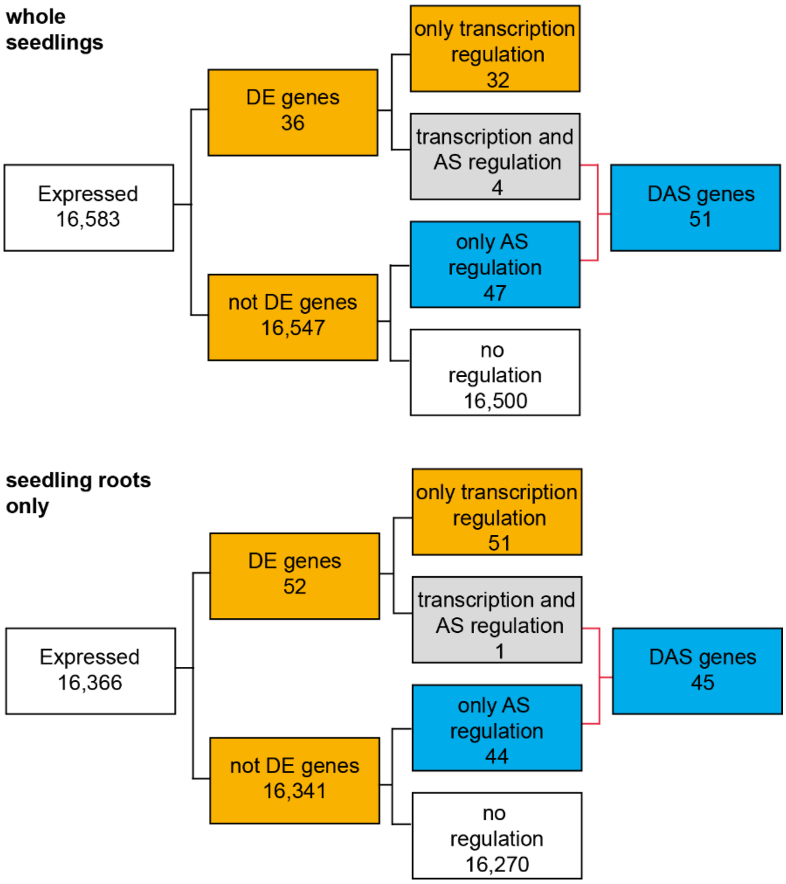


**Figure S6: RBP45 misexpression causes relatively few changes in AS and gene expression.** Numbers of expressed, differentially expressed (DE), and differentially alternatively spliced (DAS) genes in whole seedling (upper) and roots only (lower) samples. DE/DAS refers to changes in at least one RBP45 misexpression line relative to WT. The following mutant lines have been included; whole seedlings: *rbp45bc*, *rbp45abc*, *OE-A*, *OE-B* and *OE-C*; roots only: *rbp45bc*, *rbp45abc* and *OE-C* lines. The 3D RNA-seq analysis was performed with the following cut-offs for DE/DAS: p-value cut-off 0.01; log2 fold change cut-off 1; and delta PS cut-off 0.1.

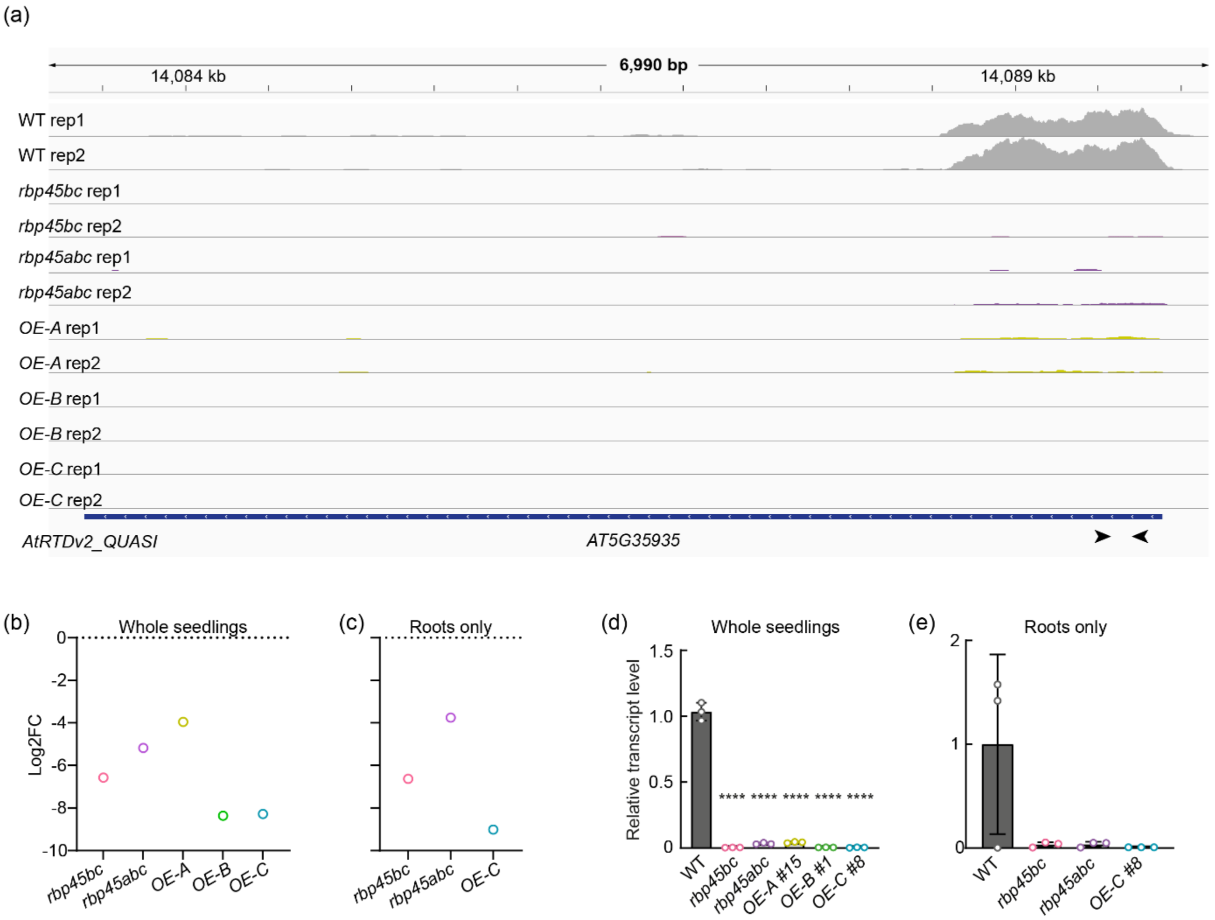


**Figure S7: A *Copia-like retrotransposon* is downregulated in all *rbp45* misexpression lines.**
(a) Coverage plots from transcriptome sequencing data of 10-d-old *A. thaliana* seedlings of indicated genotypes; all scaled to maximum of 149 reads per position. AtRTDv2_QUASI served as reference genome; the gene model of the *copia-like retrotransposon AT5G35935* is displayed at the bottom with arrows depicting primer binding sites for RT-qPCR analysis shown in (d, e).
(b, c) Change in *AT5G35935* transcript levels in comparison of the RNA-seq data from WT and *rbp45* misexpression lines. The values are negative and indicate downregulation in the mutants relative to WT. Data points represent mean values of two (b) or three (c) biological replicates.
(d, e) RT-qPCR analysis of total transcript level of *AT5G35935* from seedling (d) or root (e) samples. All values are expressed relative to *PP2A* and normalized to the respective mean WT. Mean values (bars), standard deviation (error bars), and individual data points (circles) are depicted. Asterisks indicate significant change compared to WT (one-way ANOVA followed by Dunnett's multiple comparisons test; **** $p < 0.0001$).

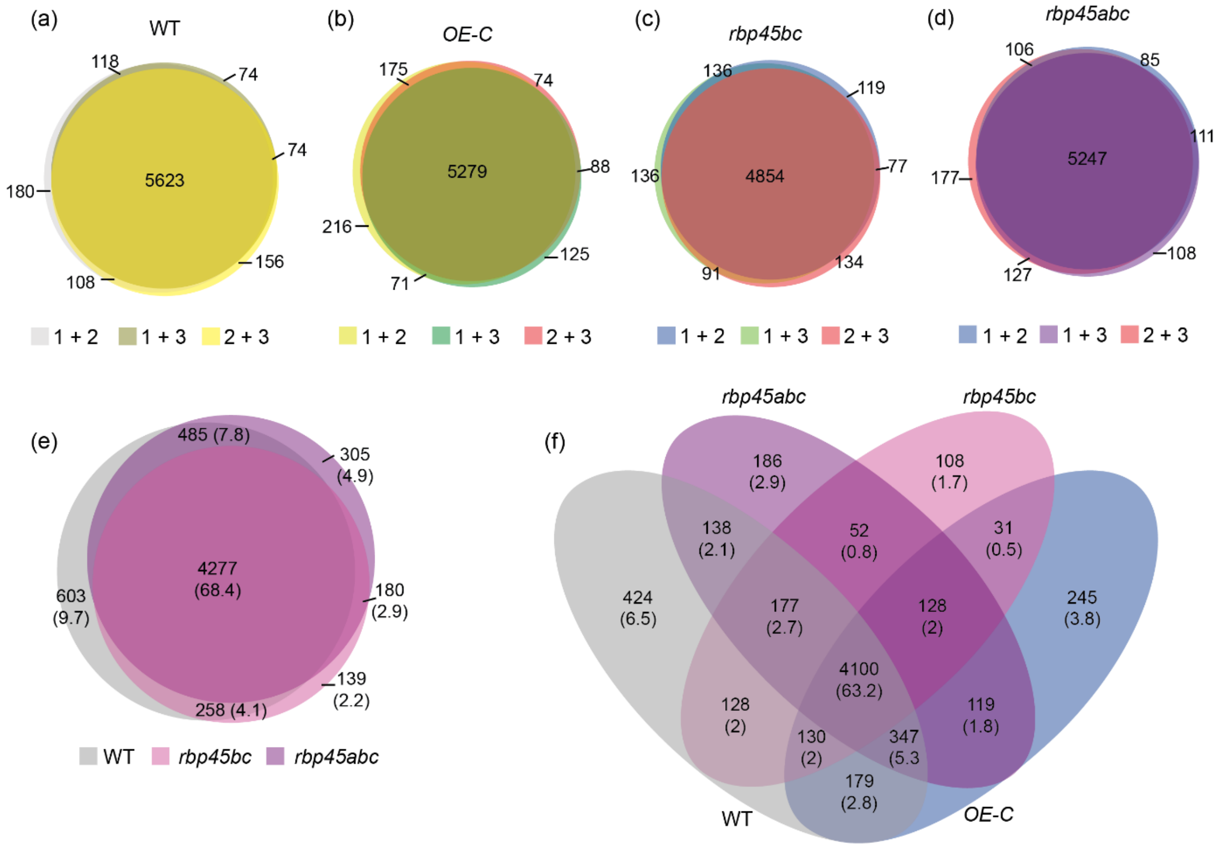


**Figure S8: Tissue-specific gene expression in WT and *rbp45* misexpression lines.**
(a – d) Comparison of the overlap in DE genes from 10-day-old seedling transcriptome and the 7-day-old root transcriptome for *A. thaliana* WT (a), *RBP45C* overexpression line (b), *rbp45bc* (c), and *rbp45abc* (d). Comparisons include different sets of replicates (indicated by numbers), as for each set of root and whole seedling samples, respectively, 3 and 2 replicates were available, and the analysis required identical number of samples for each sample type. Values in and next to circles depict the number of genes, with numbers in parenthesis providing the percentages.
(e, f) Combination of genotype-specific comparisons from (a -d) of WT and the two *rbp45* knockout lines (d) or in addition the overexpression line *OE-C* (f). For each genotype, only DE genes detected in all replicates were included.

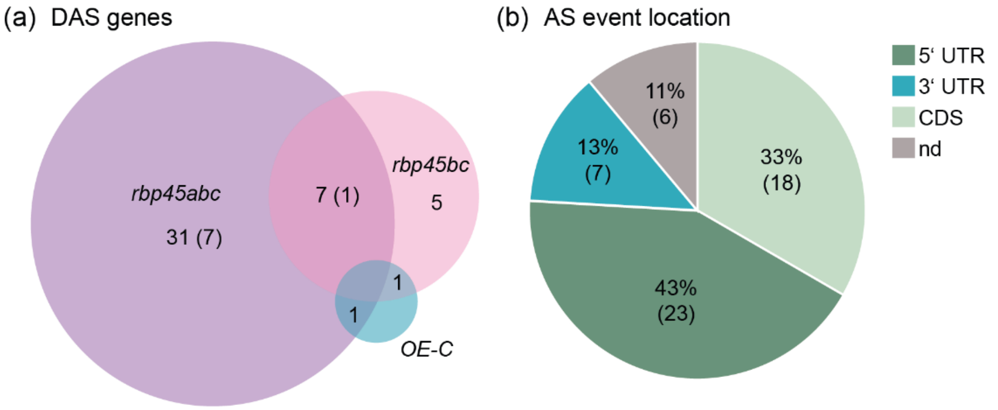


(c)

| Location assignment | Number of events |
|---|---|
| 5’ UTR events | 3,359 |
| CDS events | 22,138 |
| 3’ UTR events | 3,035 |
| 5’ UTR + CDS events | 411 |
| CDS + 3’ UTR events | 597 |
| 5’ UTR + CDS + 3’ UTR events | 32 |
| Not localized events | 22 |

**Figure S9: DAS genes and AS event location in root samples of RBP45 misexpression lines.**
(a) Venn diagram showing numbers of all significant DAS genes in root samples of 7-day-old *rbp45bc*, *rbp45abc*, and *OE-C* mutants compared to WT. Numbers in parenthesis indicate overlapping genes with seedling data set displayed in Fig. 8d.
(b) Pie chart showing for root samples location of *RBP45*-dependent AS events within the transcripts in proportions and total numbers (in parentheses). For further information and distribution of all events see Fig. 8e.
(c) Location assignments for all events based on AtRTD2-QUASI. Note that the following categories were combined under “nd” in Fig. 8e: 5’ UTR + CDS events; CDS + 3’ UTR events; 5’ UTR + CDS + 3’ UTR events; Not localized events. Additional 8,175 events could not be assigned to any of the basic AS types and therefore were not considered for the location analysis.

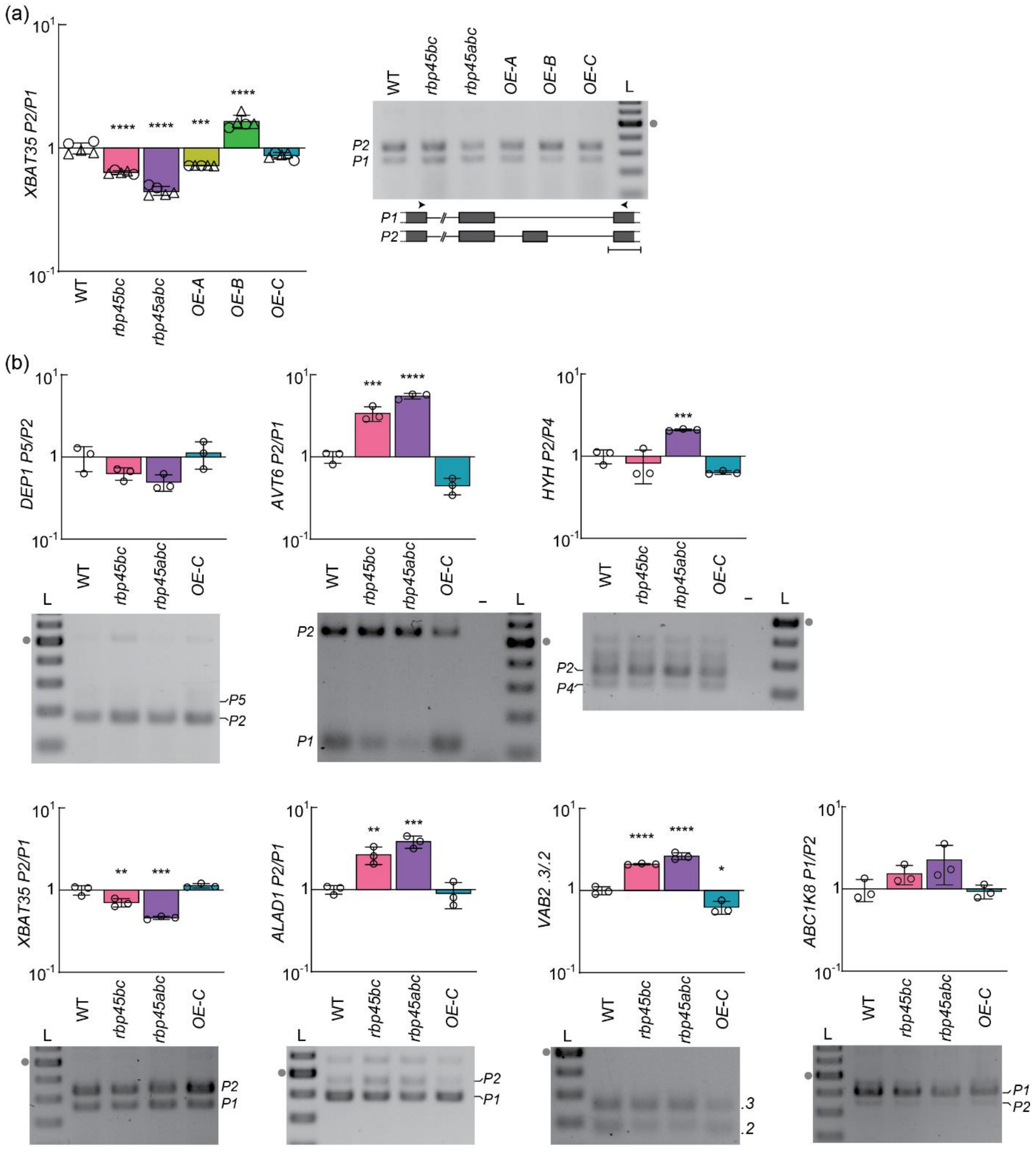


**Figure S10. Validation of splicing patterns for candidate genes in RBP45 knockout and overexpression lines.**

AS analysis from whole 10-day-old seedlings (a) or roots of 7-day-old seedlings (b) for indicated genes. Bar charts show variant ratios based on Bioanalyzer quantification of co-amplification RT-PCR products. Mean value (bars), standard deviation (error bars), and individual data points (circles: based on samples used for RNA sequencing; triangles: additional replicates) are depicted each; mean AS ratio of WT was set to 1. Asterisks indicate significant change compared to WT (one-way ANOVA followed by Dunnett's multiple comparisons test, $*p < 0.05$, $**p < 0.01$, $***p < 0.001$, $****p < 0.0001$). Representative gel pictures are shown next to or below the bar charts. Models of *XBAT35* splicing variants are shown below the gel in (a) with scale bar corresponding to 100 bp; models for other genes are displayed in Figure 9 and elements are defined in the respective legend. Size ladder (L) for gels consisted of DNAs in 100 bp increments with the strongest band corresponding to 500 bp (marked with gray dot).

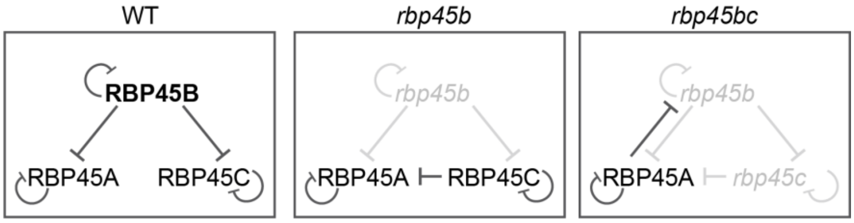


**Figure S11. Model of crosstalk and hierarchy in RBP45-mediated splicing regulation.** Negative feedback regulation in *A. thaliana* WT, and the knockout mutants *rbp45b* and r*bp45bc*.

**Supplemental Methods**

**Read trimming**

BBDuk (v39.38) was executed with the following parameters:

ref=/bbmap/resources/adapters.fa ktrim=r k=23 mink=11 hdist=1 qtrim=r trimq=20 maq=20 ml=20 tpe tbo

**Salmon quantification**

For the Salmon (v1.10.2) quantification a Salmon index was created from AtRTDv2_QUASI_19April2016.fa and the Arabidopsis genome (TAIR10_chr_all.fas) as decoy sequence.

**3D RNA-seq settings**

The following settings were used:

Data generation:

- Step 3: tximport method: lengthScaledTPM

Data pre-processing:

- Step 1: CPM cut-off: 4, Sample number cut-off: 1
- Step 4: Data normalization: TMM

3D analysis:

- Step 3: limma-voom pipeline; P-value adjust method: BH; Summarise to DAS gene level p-value: F-test; Adjusted p-value: 0.01; Absolute log_2 FC: 1; Absolute delta PS: 0.1

**Determination of AS event locations**

- determination of longest ORF per gene from FASTA file of AtRTDv2_QUASI transcriptome (AtRTDv2_QUASI_19April2016.fa)
- extraction of exon coordinates of transcripts from GTF file of AtRTDv2_QUASI transcriptome (AtRTDv2_QUASI_19April2016.gtf)
- transcript with longest ORF set as reference
- calculation of translation start and end position of reference transcript
- check whether first detectable AS event of a transcript isoform locates in 5' UTR, CDS, 3' UTR, or in overlapping regions
- only count AS event, if it is new for the current gene (= deviating coordinate is different from previous deviating coordinates for a given AS event type)
- detection of exitron: end coordinate of exons unequal; end coordinate of reference > end coordinate of isoform; end coordinate of current exon of reference coincides with end coordinate of following exon of isoform
- detection of alternative donor site: end coordinate of exons unequal; start coordinate of following exon of reference coincides with start coordinate of following exon of isoform
- detection of alternative acceptor site: start coordinate of exons unequal; end coordinate of current exon of reference coincides with end coordinate of current exon of isoform
- detection of exon skipping: start coordinate of exons unequal; start coordinate of reference < start coordinate of isoform; start coordinate of the following exon of reference coincides with start coordinate of current exon of isoform

- detection of exon insertion: start coordinates of currently examined exons of reference and isoform are unequal; start coordinate of reference > start coordinate of isoform; start coordinate of current exon of reference coincides with start coordinate of the following exon of isoform
- detection of intron retention: end coordinate of exons unequal; end coordinate of reference < end coordinate of isoform; end coordinate of following exon of reference coincides with end coordinate of current exon of isoform
- detection of mutually exclusive exons: start coordinate of exons unequal; current exon of isoform located in front of current exon of reference or behind it and start coordinates of the following exons of reference and isoform coincide

### Outcome of analysis

examined genes: 34212

proportion of examined genes for which no AS event check took place: 0

examined transcripts: 82190

Intron retention events: {'5UTR': 1031, 'CDS': 7808, '3UTR': 985, 'not localized': 0}

Alternative acceptor site events: {'5UTR': 552, 'CDS': 5339, '3UTR': 842, 'not localized': 0}

Alternative donor site events: {'5UTR': 977, 'CDS': 5438, '3UTR': 665, 'not localized': 0}

Exon insertion events: {'5UTR': 35, 'CDS': 252, '3UTR': 54, 'not localized': 0}

Exon skipping events: {'5UTR': 51, 'CDS': 848, '3UTR': 60, '5UTR+CDS': 49, 'CDS+3UTR': 53, '5UTR+CDS+3UTR': 0, 'not localized': 0}

Exitron events: {'5UTR': 713, 'CDS': 2453, '3UTR': 429, '5UTR+CDS': 362, 'CDS+3UTR': 544, '5UTR+CDS+3UTR': 32, 'not localized': 0}

Mutually exclusive exons events: {'not localized': 22}

Duplicate intron retention events: 2567

Duplicate alternative acceptor site events: 2061

Duplicate alternative donor site events: 2853

Duplicate exon insertion events: 153

Duplicate exon skipping events: 437

Duplicate exitron events: 2126

Sum of intron retention events: 12391

Sum of alternative acceptor site events: 8794

Sum of alternative donor site events: 9933

Sum of exon insertion events: 494

Sum of exon skipping events: 1498

Sum of exitron events: 6659

Sum of unresolved events: 8175

all 5UTR events: 3359

all CDS events: 22138

all 3UTR events: 3035

all 5UTR+CDS events: 411

all CDS+3UTR events: 597

all 5UTR+CDS+3UTR events: 32

all not localized events: 22

### Python script for determination of longest ORFs per gene

```
import csv, re, sys

from Bio import SeqIO


usage = "Usage: " + sys.argv[0] + " <path/transcriptome file (fa file)>" + " <path/longest orf file (txt file)>"

# transcriptome file: contains transcript sequences, from which longest orfs have to be retrieved

# longest orf file: filename of file for export of longest orfs per gen


if len(sys.argv) != 3:
  print(len(sys.argv))
  print(usage)
  sys.exit()


def find_longest_ORF4transcript(RNA):
   ORFs = []
   if 'AUG' in RNA:
     for startMatch in re.finditer('AUG', RNA):
       remaining = RNA[startMatch.start():]
       for stopMatch in re.finditer('UAA|UGA|UAG', remaining):
         substring = remaining[:stopMatch.end()]
         if len(substring) % 3 == 0:
           ORFs.append(substring)
           # first stopcodon alone relevant because translation ends here
           break
   ORFs.sort(key=len, reverse=True)
   # avoid error at return if list kept empty
```

```
  if len(ORFs) == 0:
    ORFs.append('')
  return ORFs[0]



filename = sys.argv[1]

sequences = SeqIO.parse(filename, 'fasta')
L = []

for record in sequences:
  L.append(record)



#1. Run:
#-------
#- find longest ORFs per gen
#- save these in Dictionary longest_orfs4genes

def save_current_data(current_gene, maxlen_orf, longest_orfs4genes):
  longest_orfs4genes[current_gene] = maxlen_orf

current_gene = ''
maxlen_orf = 0
longest_orfs4genes = {}

for record_ in L:
  RNA = str(record_.seq.transcribe())
  ORF = find_longest_ORF4transcript(RNA)
  if ORF != '':
    # when switching to a new gene...
    if record_.id[0:9] != current_gene:
      save_current_data(current_gene, maxlen_orf, longest_orfs4genes)
      # ... save its name in current_gene
      current_gene = record_.id[0:9]
      # ... reset maxlen
      maxlen_orf = 0
    if len(ORF) > maxlen_orf:
      maxlen_orf = len(ORF)
```

```
#2. Run:
#-------
#- build list records_ with transcript isoforms, which contain longest ORFs per accompanying gen

def save_possibly_current_record(current_gene, longest_orfs4genes, ORF, record_, records_):
  if current_gene != '':
    try:
      current_longest_orf_length4gen = longest_orfs4genes[current_gene]
    except:
      print('Laengster ORF zu Gen ', current_gene, ' existiert nicht.')
    else:
      if current_longest_orf_length4gen == len(ORF):
        records_.append(record_)

current_gene = ''
records_ = []

for record_ in L:
  RNA = str(record_.seq.transcribe())
  ORF = find_longest_ORF4transcript(RNA)
  if ORF != '':
    save_possibly_current_record(record_.id[0:9], longest_orfs4genes, ORF, record_, records_)



#Export list of records with longest ORFs per gene along with start and end coordinate and length of ORF (1-based coordinates);
#if several transcripts of a gene have equally long longest ORFs, then export only one (here: the last scrutinized):
#-----------------------------------------------------------------------------------------------------------------------------
FileHandle1 = open(sys.argv[2], 'w')

writer1 = csv.writer(FileHandle1, delimiter=',')
current_gene = ''
prev_element = object()
i = 0
for element in records_:
  if element.id[0:9] != current_gene:
    current_gene = element.id[0:9]
    # export data of sole or of last scrutinized transcript with longest orf per gene
```

```
    if i > 0:
      RNA = str(prev_element.seq.transcribe())
      ORF = find_longest_ORF4transcript(RNA)
      for match in re.finditer(ORF, RNA):
        row_ = []
        row_.append(prev_element.id)
        row_.append(match.start()+1)
        row_.append(match.end())
        row_.append(len(ORF))
        row_.append(RNA[match.start():match.end()])
        writer1.writerow(row_)
      FileHandle1.flush()
  prev_element = element
  i += 1
```

Python script for extraction of exon coordinates of transcripts:

```
import sys, re, pickle


usage = "Usage: " + sys.argv[0] + " <path/gtf file>" + " <path/exon file (pkl file)>"


if len(sys.argv) != 3:
  print(len(sys.argv))
  print(usage)
  sys.exit()



def read_gtf_file(gtf_file):
# read data from gtf file; extract transcript-ID, exon start, exon end and strand; save extracted data as tuple and put tuple into dictionary
# check, whether exons are really separated by introns; otherwise unite exons into one
# return dictionary with exon data
  data = {}
  GTF = open(gtf_file)
  lines_in_gtf = 0
  exons = 0
  exon = ()
  previous_exon = ()
  transcript_id = ''
  previous_transcript_id = ''
```

```
  transcript_counter = 0
  incorporation_counter = 0
  for line in GTF:
    previous_transcript_id = transcript_id
    previous_exon = exon
    lines_in_gtf += 1
    chrom,source,seqtype,start,end,score,strand,frame,attr = line.strip().split('\t')
    if seqtype == 'exon':
      exons += 1
      if "transcript_id" in attr:
        match = re.search('transcript_id "(.*?)";',attr)
        transcript_id = match.group(1)
        if transcript_id not in data:
          data[transcript_id] = []
          transcript_counter += 1
        # gtf file coordinates: 1-based + inclusive - let it so
        exon = (chrom, int(start), int(end), strand)
        if previous_exon != () and previous_transcript_id == transcript_id and previous_exon[2]+1 == exon[1]:
          # previous and current exon belong to the same transcript and are seamlessly connected; therefore unite them
          exon[1] = previous_exon[1]
          # don't add to dictionary, as long as furhter exons could be incorporated
          incorporation_counter += 1
        else:
          data[transcript_id].append(exon)
      else:
        print("error parsing GTF")
        exit()
  print("lines_in_gtf: ",lines_in_gtf,", Exons: ", exons, ", Transcripts: ", transcript_counter, ", Exon unitings: ", incorporation_counter)
  return data


# save exon data in a pkl file:
transcriptModels = read_gtf_file(sys.argv[1])
with open(sys.argv[2], 'wb') as fp:
    pickle.dump(transcriptModels, fp)
```

### Python script for AS event location:

```
import sys, pickle
```

```
usage = "Usage: " + sys.argv[0] + " <path/longest orf file (txt file)>" + " <path/exon file (pkl file)>"

# longest orf file: transcripts that have longest orfs among all transcripts of a gene; if several transcripts with longest orfs exist,

# one is arbitrarily choosen (= serves as reference)

# exon file: contains exon coordinates to transcripts


if len(sys.argv) != 3:
  print(len(sys.argv))
  print(usage)
  sys.exit()


def read_longest_orf_file(orf_file):
  data = []
  tx_tupel = ()
  orfFile = open(orf_file)
  for line in orfFile:
    # caution: file contains empty lines
    if line not in ['\n','\r\n']:
      try:
        tx_id,start,end,length,seq = line.strip().split(",")
        tx_tupel = (tx_id,start,end,length,seq)
        data.append(tx_tupel)
      except:
        print('read_orf_file1: Error in file reading')
  return data


def check_for_intron_insertion(transcriptModels, ref_key, isoform_key, i):
  # assumed as given: 1) end coordinates of currently examined exons of reference and isoform are unequal; 2) end coordinate of reference > end coordinate of isoform
  # check whether end coordinate of current exon of reference coincides with end coordinate of following exon of isoform
  # in this case intron insertion has happened; then
  # return raw end and start coordinates of intron-preceding and intron-following exons, respectively, otherwise empty tuple
  # limitation: in case of some complex as event (for example: next exon changed by alternative donor site), intron insertion is not recognized
  # transcriptModels: Dictionary with entries consisting of transcript ids as keys and a list of exon related tupels as value
  # ref_key = key of the reference transcript
  # isoform_key = key of the transcript isoform for comparison with the reference
  # i = index of the exon list
  ii_coordinates = ()
```

```
  try:
    if len(transcriptModels[isoform_key]) <= (i+1):
      # following comparison not possible
      print('Failure in check_for_intron_insertion(): no following exon in isoform transcript existent')
      return ii_coordinates
    if transcriptModels[ref_key][i][2] == transcriptModels[isoform_key][i+1][2]:
      print('End of raw exon ', i, ' of reference = end of raw exon ', i+1, ' of isoform: ', transcriptModels[ref_key][i][2])
      ii_coordinates = (transcriptModels[isoform_key][i][2], transcriptModels[isoform_key][i+1][1])
      return ii_coordinates
    else:
      print('no intron insertion')
  except IndexError:
    print('IndexError in check_for_intron_insertion()')
  except:
    print('Error in check_for_intron_insertion()')
  return ii_coordinates


def check_for_intron_retention(transcriptModels, ref_key, isoform_key, i):
  # assumed as given: 1) end coordinates of currently examined exons of reference and isoform are unequal; 2) end coordinate of reference < end coordinate of isoform
  # check whether end coordinate of following exon of reference coincides with end coordinate of current exon of isoform
  # in this case intron retention has happened; then
  # return true, otherwise false
  # limitation: in case of some complex as event (for example: next intron also retained), intron retention is not recognized
  # transcriptModels: Dictionary with entries consisting of transcript ids as keys and a list of exon related tupels as value
  # ref_key = key of the reference transcript
  # isoform_key = key of the transcript isoform for comparison with the reference
  # i = index of the exon list
  try:
    if len(transcriptModels[ref_key]) <= i+1:
      # no following exon in reference transcript existent, so that following comparison not possible
      print('Failure in check_for_intron_retention(): no check for ir possible')
      return False
    if transcriptModels[ref_key][i+1][2] == transcriptModels[isoform_key][i][2]:
      print('End of raw exon ', i+1, ' of reference = end of raw exon ', i, ' of isoform: ', transcriptModels[ref_key][i+1][2])
      return True
    else:
      print('no intron retention')
```

```
  except IndexError:
    print('IndexError in check_for_intron_retention()')
  except:
    print('Error in check_for_intron_retention()')
  return False


def check_for_alt_acceptor(transcriptModels, ref_key, isoform_key, i):
  # assumed as given: start coordinates of currently examined exons of reference and isoform are unequal
  # check whether end coordinate of current exon of reference coincides with end coordinate of current exon of isoform
  # limitation: in case of some complex as event (for example: next intron retained or alternative donor used in the current exon),
  # alternative acceptor event is not recognized
  raw_start_coordinate_modified_exon = ()
  try:
    if transcriptModels[ref_key][i][2] == transcriptModels[isoform_key][i][2]:
      print('End of raw exon ', i, ' of reference = end of raw exon ', i, ' of isoform: ', transcriptModels[ref_key][i][2])
      raw_start_coordinate_modified_exon = (transcriptModels[isoform_key][i][1])
      return raw_start_coordinate_modified_exon
    else:
      print('no alternative acceptor')
  except IndexError:
    print('IndexError in check_for_alt_acceptor()')
  except:
    print('Error in check_for_alt_acceptor()')
  return raw_start_coordinate_modified_exon


def check_for_alt_donor(transcriptModels, ref_key, isoform_key, i):
  # assumed as given: end coordinates of currently examined exons of reference and isoform are unequal
  # check whether start coordinate of following exon of reference coincides with start coordinate of following exon of isoform
  raw_end_coordinate_modified_exon = ()
  try:
    if len(transcriptModels[ref_key]) <= i+1 or len(transcriptModels[isoform_key]) <= i+1:
      # no following exon in reference or isoform transcript existent, so that following comparison not possible
      print('Failure in check_for_alt_donor(): no check for altd possible')
      return raw_end_coordinate_modified_exon
    if transcriptModels[ref_key][i+1][1] == transcriptModels[isoform_key][i+1][1]:
         print('Start of raw exon of reference ', i+1, ' = start of raw exon of isoform ', i+1, ' : ', transcriptModels[ref_key][i+1][1])
      raw_end_coordinate_modified_exon = (transcriptModels[isoform_key][i][2])
```

```
      return raw_end_coordinate_modified_exon
  except IndexError:
    print('IndexError in check_for_alt_donor()')
  except:
    print('Error in check_for_alt_donor()')
  return raw_end_coordinate_modified_exon


def check_for_exon_insertion(transcriptModels, ref_key, isoform_key, i):
  # assumed as given: 1) start coordinates of currently examined exons of reference and isoform are unequal; 2) start coordinate of reference > start coordinate of isoform
  # check whether start coordinate of current exon of reference coincides with start coordinate of the following exon of isoform
  try:
    if len(transcriptModels[isoform_key]) <= i+1:
      # no following exon in isoform transcript existent, so that following comparison not possible
      print('Failure in check_for_exon_insertion(): no check for exonin possible')
      return False
    if transcriptModels[ref_key][i][1] == transcriptModels[isoform_key][i+1][1]:
      print('Start of raw exon of reference ', i, ' = start of raw exon of isoform ', i+1, ' : ', transcriptModels[ref_key][i][1])
      return True
  except IndexError:
    print('IndexError in check_for_exon_insertion()')
  except:
    print('Error in check_for_exon_insertion()')
  return False


def check_for_exon_skipping(transcriptModels, ref_key, isoform_key, i):
  # assumed as given: 1) start coordinates of currently examined exons of reference and isoform are unequal; 2) start coordinate of reference < start coordinate of isoform
  # check whether start coordinate of the following exon of reference coincides with start coordinate of current exon of isoform
  # in this case exon_skipping has happened; then return start and end coordinates of skipped exon + end coordinate of previous exon +
  # start coordinate of next exon, otherwise empty tuple
  exonskip_coordinates = ()
  try:
    if len(transcriptModels[ref_key]) <= i+1:
      # no following exon in reference transcript existent, so that following comparison not possible
      print('Failure in check_for_exon_skipping(): no following exon in reference transcript existent')
      return exonskip_coordinates
    if transcriptModels[ref_key][i+1][1] == transcriptModels[isoform_key][i][1]:
```

```
    print('Start of raw exon of reference ', i+1, ' = start of raw exon of isoform ', i, ' : ', transcriptModels[ref_key][i+1][1])
    exonskip_coordinates = (transcriptModels[ref_key][i][1], transcriptModels[ref_key][i][2], transcriptModels[ref_key][i-1][2], transcriptModels[ref_key][i+1][1])
    return exonskip_coordinates
  except IndexError:
    print('IndexError in check_for_exon_skipping()')
  except:
    print('Error in check_for_exon_skipping()')
  return exonskip_coordinates


def check_for_mutually_exclusive_exons(transcriptModels, ref_key, isoform_key, i):
  # assumed as given: 1) start coordinates of currently examined exons of reference and isoform are unequal
  # check whether the current exon of isoform is located in front of the current exon of reference or behind it and
  # check whether start coordinates of the following exons of reference and isoform coincide
  # in this case exon_skipping has happened; then return start and end coordinates of skipped exon + end coordinate of previous exon +
  # start coordinate of next exon, otherwise empty tuple
  #exonskip_coordinates = ()
  try:
    if len(transcriptModels[ref_key]) <= i+1:
      # no following exon in reference transcript existent, so that following comparison not possible
      print('Failure in check_for_exon_skipping(): no following exon in reference transcript existent')
      return exonskip_coordinates
    if (transcriptModels[isoform_key][i][2] < transcriptModels[ref_key][i][1] or transcriptModels[isoform_key][i][1] > transcriptModels[ref_key][i][2]) and (transcriptModels[ref_key][i+1][1] == transcriptModels[isoform_key][i+1][1]):
      print('Start of raw exon of reference ', i+1, ' = start of raw exon of isoform ', i+1, ' : ', transcriptModels[ref_key][i+1][1])
      exonskip_coordinates = (transcriptModels[ref_key][i][1], transcriptModels[ref_key][i][2], transcriptModels[ref_key][i-1][2], transcriptModels[ref_key][i+1][1])
      #return exonskip_coordinates
      return True
  except IndexError:
    print('IndexError in check_for_mutually_exclusive_exons()')
  except:
    print('Error in check_for_mutually_exclusive_exons()')
  #return exonskip_coordinates
  return False


def detect_as_event(transcriptModels, ref_key, isoform_key, ir_list, alta_list, altd_list, exonin_list, ii_list, unresolved_list, exonskip_list):
  # compare exon boundaries of reference transcript and isoform in parallel; assign first deviation to an as event
```

```
  # returns tupel containing first deviation coordinate and as event code (ir = 1, alta = 2, altd = 3, exonin = 4, ii = 5, exonskip = 6, mutexexon = 7) or tuple (0, 0) for unresolved event

  # parameters:

  # transcriptModels = dictionary containing exon data for transcripts originating from gtf file (transcript ids as keys and a list of exon related tupels as value)

  # ref_key = key of the reference transcript

  # isoform_key = key of the transcript isoform for comparison with the reference

  # ir_list = list of ir Events, containing Tupels of transcript_id + number of retained intron - to be further filled here

  # alta_list = list of alternative Acceptor Events, containing Tupels of transcript_id + number of affected exon - to be further filled here

  as_event = ()

  deviating_exon_coordinate = 0

  as_event_code = 0

  i = 0

  ii_t = ()

  altd_t = ()

  alta_t = ()

  exonskip_t = ()

  #isoform_value = transcriptModels[isoform_key]

  for ref_entry in transcriptModels[ref_key]:

    # for loop picks up exons of reference transcript

    try:

      if ref_entry[1] != transcriptModels[isoform_key][i][1]:

            print('Deviation in raw start coordinate of exon ', i, ': ', ref_entry[1], '(reference) != ', transcriptModels[isoform_key][i][1], ' (isoform)')

        deviating_exon_coordinate = transcriptModels[isoform_key][i][1]

        # mutually exclusive exon

        if check_for_mutually_exclusive_exons(transcriptModels, ref_key, isoform_key, i) == True:

          print('mutually_exclusive_exons')

          as_event_code = 7

          break

        # alternative acceptor: isoform exon can be either shorter (starts later) or longer (starts earlier) than reference exon

        alta_t = check_for_alt_acceptor(transcriptModels, ref_key, isoform_key, i)

        if alta_t != ():

          print('alternative acceptor')

          # save transcript id of isoform and coordinate of modified exon:

          alta_list.append((isoform_key, alta_t))

          as_event_code = 2

          # treat only first coordinate deviation

          break
```

```
        # exon insertion: reference exon starts later than (inserted) exon of isoform
        if ref_entry[1] > transcriptModels[isoform_key][i][1]:
          if check_for_exon_insertion(transcriptModels, ref_key, isoform_key, i) == True:
            print('exon insertion')
            # save transcript id of isoform and number of previous intron (where additional exon of isoform is inserted):
            exonin_list.append((isoform_key, i-1))
            as_event_code = 4
            break
        # exon skipping: reference exon starts earlier than exon of isoform, which follows on not more present exon
        if ref_entry[1] < transcriptModels[isoform_key][i][1]:
            exonskip_t = check_for_exon_skipping(transcriptModels, ref_key, isoform_key, i)
            if exonskip_t != ():
              print('exon skipping')
               # save transcript id of isoform and coordinates of skipped exon of reference and coordinates of adjacent exons:
              exonskip_list.append((isoform_key, exonskip_t))
              as_event_code = 6
              break
        print('unresolved as event')
        unresolved_list.append((isoform_key, i))
        # treat only first coordinate deviation
        break
      if ref_entry[2] != transcriptModels[isoform_key][i][2]:
            print('Deviation in raw end coordinate of raw exon ', i, ': ', ref_entry[2], '(reference) != ', transcriptModels[isoform_key][i][2], ' (isoform)')
        deviating_exon_coordinate = transcriptModels[isoform_key][i][2]
        # alternative donor: isoform exon can be either shorter (ends earlier) or longer (ends later) than reference exon
        altd_t = check_for_alt_donor(transcriptModels, ref_key, isoform_key, i)
        if altd_t != ():
          print('alternative donor')
          # save transcript id of isoform and coordinate of modified exon:
          altd_list.append((isoform_key, altd_t))
          as_event_code = 3
          break
        # intron insertion: reference exon longer than intron-preceding exon of isoform
        if ref_entry[2] > transcriptModels[isoform_key][i][2]:
          ii_t = check_for_intron_insertion(transcriptModels, ref_key, isoform_key, i)
          if ii_t != ():
            print('exitron')
            # save transcript id of isoform and coordinates of adjacent exons:
```

```
          ii_list.append((isoform_key, ii_t))
          as_event_code = 5
          break
      # intron retention: reference exon shorter than isoform exon (which contains former intron)
      if ref_entry[2] < transcriptModels[isoform_key][i][2]:
        if check_for_intron_retention(transcriptModels, ref_key, isoform_key, i) == True:
          print('retained intron')
          # save transcript id of isoform and number of retained intron:
          ir_list.append((isoform_key, i))
          as_event_code = 1
          break
      print('unresolved as event')
      unresolved_list.append((isoform_key, i))
      break
    except IndexError:
      print('IndexError')
      break
    except:
      print('Error other than IndexError')
      break
    i += 1
  as_event = (deviating_exon_coordinate, as_event_code)
  return as_event


def get_intronless_translation_coordinates(ref_transcript_id, transcriptModels, orf_data):
# Returns translation start and end coordinates for handed over reference transcript
# ref_transcript_id: transcript id of reference transcript
# transcriptModels = dictionary containing exon data for transcripts originating from gtf file of transcriptome (transcript ids as keys and a list of exon related tupels as value)
# orf_data = list of tupels containing reference transcript id, start coordinate of orf (= translation start),
# end coordinate of orf (= translation end + 3), length of orf, sequence of orf; all these extracted from fa file of transcriptome
  exon_start = 0
  offset_start = 0
  offset_end = 0
  translation_coordinates = ()
  for key, value in transcriptModels.items():
    if key == ref_transcript_id:
      # gtf coordinate (1-based, inclusive) already converted into python coordinate (0-based, exclusive)
      exon_start = value[0][1]
```

```
    for entry in orf_data:
      #print('get_intronless_translation_coordinates(): ', entry)
      if entry[0] == key:
        # determine translation coordinates for reference transkript
        offset_start = int(entry[1])
        # offset_end calculation: minus 3 because orf contains stopcodon
        offset_end = int(entry[2])-3
        translation_coordinates = (ref_transcript_id, exon_start + offset_start - 1, exon_start + offset_end - 1)
        break
    break
  return translation_coordinates


def get_intronless_exon_coordinates(transcript_id, transcriptModels):
# Returns list of exon coordinates for handed over transcript id after subtraction of introns;
# list entries: Tupels containing transcript_id, chromosome, exon start, exon end, accumulated intron length until current exon, strand
# transcript_id: transcript id
# transcriptModels = dictionary containing exon data for transcripts originating from gtf file (transcript ids as keys and a list of exon related tupels as value)
  exons = []
  accumulating_intron_length = 0
  for key, value in transcriptModels.items():
    if key == transcript_id:
      i = 0
      # first exon needs no change
      exons.append((transcript_id, value[i][0], value[i][1], value[i][2], 0, value[i][3]))
      # treat all further exons of this transcript:
      while i < len(value)-1:
        # Intron length of intron i: (start coordinate of exon i+1) - (end coordinate of exon i) - 1
        # (subtraction of 1, because coordinates of exons don't belong to intron)
        accumulating_intron_length += value[i+1][1] - value[i][2] - 1
        exons.append((transcript_id, value[i+1][0], value[i+1][1]-accumulating_intron_length, value[i+1][2]-accumulating_intron_length, accumulating_intron_length, value[i+1][3]))
        i += 1
      break
  return exons


def get_intronless_translation_data(ref_transcript_id, transcriptModels, orf_data, intronless_exons):
# Returns Tuple containing handed over reference transcript id, accompanying start coordinate of translation, stop coordinate of translation,
#  start exon of translation, stop exon of translation, assignment status of start exon, assignment status of stop exon
```

```
# ref_transcript_id: transcript id of reference transcript

# transcriptModels = dictionary containing exon data for transcripts originating from gtf file (transcript ids as keys and a list of exon related tupels as value)

# orf_data = list of tupels containing reference transcript id, start coordinate of orf, end coordinate of orf, length of orf, sequence of orf

# intronless_exons = list of exon coordinates for reference transcript after subtraction of introns; list entries: Tupels containing transcript_id, chromosome,

#  exon start, exon end, accumulated intron length until current exon, strand

  intronless_translation_coordinates = ()

  translation_data = ()

  start_exon_number = 0

  stop_exon_number = 0

  start_exon_assigned = False

  stop_exon_assigned = False

  i = 0

  try:

      intronless_translation_coordinates = get_intronless_translation_coordinates(ref_transcript_id, transcriptModels, orf_data)

    for exon in intronless_exons:

      if (intronless_translation_coordinates[1] >= exon[2]) and (intronless_translation_coordinates[1] <= exon[3]):

        start_exon_number = i

        start_exon_assigned = True

      if (intronless_translation_coordinates[2] >= exon[2]) and (intronless_translation_coordinates[2] <= exon[3]):

        stop_exon_number = i

        stop_exon_assigned = True

      i += 1

     translation_data = (ref_transcript_id, intronless_translation_coordinates[1], intronless_translation_coordinates[2], start_exon_number, stop_exon_number, start_exon_assigned, stop_exon_assigned)

  except IndexError:

    print('IndexError in get_intronless_translation_data()')

  except:

    print('Error in get_intronless_translation_data()')

  return translation_data


def get_raw_translation_coordinates(ref_transcript_id, transcriptModels, orf_data, intronless_exons):

# Returns tuple containing raw start and end coordinate of translation for the transcript with handed over reference transcript id

# ref_transcript_id: transcript id of reference transcript

# transcriptModels = dictionary containing exon data for transcripts originating from gtf file (transcript ids as keys and a list of exon related tupels as value)

# orf_data = list of tupels containing reference transcript id, start coordinate of orf, end coordinate of orf, length of orf, sequence of orf
```

```
# intronless_exons = list of exon coordinates for reference transcript after subtraction of introns; list entries: Tupels containing transcript_id, chromosome,
#  exon start, exon end, accumulated intron length until current exon, strand
  intronless_translation_data = ()
  raw_translation_start_coordinate = 0
  raw_translation_end_coordinate = 0
  i = 0
  intronless_translation_data = get_intronless_translation_data(ref_transcript_id, transcriptModels, orf_data, intronless_exons)
  for i_exon in intronless_exons:
    if i_exon[0] == ref_transcript_id and i == intronless_translation_data[3]:
      # just scrutinized exon is start exon of translation -> add accumulated intron length until this exon
      raw_translation_start_coordinate = intronless_translation_data[1] + i_exon[4]
    if i_exon[0] == ref_transcript_id and i == intronless_translation_data[4]:
      # just scrutinized exon is end exon of translation -> add accumulated intron length until this exon
      raw_translation_end_coordinate = intronless_translation_data[2] + i_exon[4]
    i += 1
  return (raw_translation_start_coordinate, raw_translation_end_coordinate)


def get_ii_localization(transcript_id, ii_list, raw_translation_coordinates):
# determine whether ii happened in 5'UTR, CDS, 3'UTR, 5'UTR+CDS, CDS+3'UTR, or 5'UTR+CDS+3'UTR
# transcript_id: transcript id of isoform to be examined
# ii_list: list of ii events, containing tuples of transcript id + tuple of exon coordinates needed for localization
# raw_translation_coordinates: Tuple containing raw start and end coordinate of translation
# Returns 0 for unsuccessful localization, 1 for 5'UTR, 2 for CDS, 3 for 3'UTR, 4 for 5'UTR+CDS, 5 for CDS+3'UTR, 6 for 5'UTR+CDS+3'UTR
  localization = 0
  raw_isoform_end_coordinate_exon_ahead_inserted_intron = 0
  raw_isoform_start_coordinate_exon_after_inserted_intron = 0

  # get raw start and end coordinate of adjacent exons:
  # - find corresponding list entry in ii_list:
  for entry in ii_list:
    if entry[0] == transcript_id:
      raw_isoform_end_coordinate_exon_ahead_inserted_intron = entry[1][0]
      raw_isoform_start_coordinate_exon_after_inserted_intron = entry[1][1]
      break

  # comparisons
  if raw_translation_coordinates[1] <= raw_isoform_end_coordinate_exon_ahead_inserted_intron:
    # event in 3'UTR
```

```
    print('* Reverse IR in transcript isoform ', transcript_id, ' happened in 3UTR')
    localization = 3
    return localization
  if raw_translation_coordinates[0] >= raw_isoform_start_coordinate_exon_after_inserted_intron:
    # event in 5'UTR
    print('* Reverse IR in transcript isoform ', transcript_id, ' happened in 5UTR')
    localization = 1
    return localization
  if raw_translation_coordinates[0] <= raw_isoform_end_coordinate_exon_ahead_inserted_intron+1 and raw_translation_coordinates[1] >= raw_isoform_start_coordinate_exon_after_inserted_intron-1:
    # event in CDS
    print('* Reverse IR in transcript isoform ', transcript_id, ' happened in CDS')
    localization = 2
    return localization
  if raw_translation_coordinates[0] > raw_isoform_end_coordinate_exon_ahead_inserted_intron+1 and raw_translation_coordinates[0] < raw_isoform_start_coordinate_exon_after_inserted_intron and raw_translation_coordinates[1] >= raw_isoform_start_coordinate_exon_after_inserted_intron-1:
    # event in 5UTR + CDS
    print('* Reverse IR in transcript isoform ', transcript_id, ' happened in 5UTR and CDS')
    localization = 4
    return localization
  if raw_translation_coordinates[0] <= raw_isoform_end_coordinate_exon_ahead_inserted_intron+1 and raw_translation_coordinates[1] > raw_isoform_end_coordinate_exon_ahead_inserted_intron and raw_translation_coordinates[1] < raw_isoform_start_coordinate_exon_after_inserted_intron-1:
    # event in CDS + 3UTR
    print('* Reverse IR in transcript isoform ', transcript_id, ' happened in CDS and 3UTR')
    localization = 5
    return localization
  if raw_translation_coordinates[0] > raw_isoform_end_coordinate_exon_ahead_inserted_intron+1 and raw_translation_coordinates[1] < raw_isoform_start_coordinate_exon_after_inserted_intron-1:
    # event in 5UTR + CDS + 3UTR
    print('* Reverse IR in transcript isoform ', transcript_id, ' happened in 5UTR and CDS and 3UTR')
    localization = 6
    return localization

  return localization

def get_ir_localization(ref_transcript_id, transcript_id, ir_list, intronless_translation_data):
# determine whether ir happened in 5'-UTR, CDS or 3'-UTR
# ref_transcript_id: transcript id of reference transcript
# transcript_id: transcript id of isoform to be examined
# ir_list: list of ir events, containing Tupels of transcript_id + number of retained intron
```

```
# intronless_translation_data: Tupel containing handed over reference transcript id, start coordinate of translation, stop coordinate of translation,
#  start exon of translation, stop exon of translation
# Returns 0 for unsuccessful localization, 1 for 5'UTR, 2 for CDS, 3 for 3'UTR
# logic:
# Input: sequence exon 0 - intron 0 - exon 1 - intron 1 - ...
# if translation start in or downstream to exon succeeding retained intron, then ir in 5'UTR
# else if translation end upstream to exon succeeding retained intron, then ir in 3'UTR
# else ir in CDS
  localization = 0
  # fish for number of retained intron
  intron_number = -1
  for entry in ir_list:
    if entry[0] == transcript_id:
      intron_number = entry[1]
      break
  print('get_ir_localization(): intron_number = ', intron_number)
  # find number of exon of reference after retained intron
  succeeding_exon_number = intron_number+1

  if intronless_translation_data[0] == ref_transcript_id and intron_number != -1:
    if intronless_translation_data[3] >= succeeding_exon_number:
      print('* IR in transcript isoform ', transcript_id, ' happened in 5UTR')
      localization = 1
    if intronless_translation_data[4] < succeeding_exon_number:
      print('* IR in transcript isoform ', transcript_id, ' happened in 3UTR')
      localization = 3
    if intronless_translation_data[3] < succeeding_exon_number and intronless_translation_data[4] >= succeeding_exon_number:
      print('* IR in transcript isoform ', transcript_id, ' happened in CDS')
      localization = 2

  return localization

def get_alta_localization(transcript_id, alta_list, raw_translation_coordinates):
# determine whether alta happened in 5'-UTR, CDS or 3'-UTR
# transcript_id: transcript id of isoform to be examined
# alta_list: list of alta events, containing Tupels of transcript_id + number of modified exon
# raw_translation_coordinates: Tuple containing raw start and end coordinate of translation
# Returns 0 for unsuccessful localization, 1 for 5'UTR, 2 for CDS, 3 for 3'UTR
```

```
# logic:
# import raw start coordinate of modified (either elongated or shortened) exon
# if modified start coordinate at or downstream translation start (ts), event in CDS or in 3'UTR, otherwise (= upstream ts) event in 5'UTR
# in case of at or downstream ts: if modified start coordinate at or upstream translation end (te), event in CDS;
#  else (= downstream te), event in 3'UTR
  localization = 0
  raw_start_coordinate_modified_exon = 0
  # get raw start coordinate of modified exon:
  for entry in alta_list:
    if entry[0] == transcript_id:
      raw_start_coordinate_modified_exon = entry[1]
      break
  # comparisons between event coordinate and translation coordinates:
  if raw_start_coordinate_modified_exon >= raw_translation_coordinates[0]:
    # event in CDS or 3UTR
    if raw_start_coordinate_modified_exon <= raw_translation_coordinates[1]:
      # event in CDS
        print('* AltA in transcript isoform ', transcript_id, ' happened in CDS - AS event at ', raw_start_coordinate_modified_exon)
      localization = 2
    else:
        print('* AltA in transcript isoform ', transcript_id, ' happened in 3UTR - AS event at ', raw_start_coordinate_modified_exon)
      localization = 3
  else:
    # event in 5UTR
      print('* AltA in transcript isoform ', transcript_id, ' happened in 5UTR - AS event at ', raw_start_coordinate_modified_exon)
    localization = 1

  return localization

def get_altd_localization(transcript_id, altd_list, raw_translation_coordinates):
# determine whether altd happened in 5'-UTR, CDS or 3'-UTR
# transcript_id: transcript id of isoform to be examined
# altd_list: list of altd events, containing Tupels of transcript_id + number of modified exon
# raw_translation_coordinates: Tupel containing raw start and end coordinate of translation
# Returns 0 for unsuccessful localization, 1 for 5'UTR, 2 for CDS, 3 for 3'UTR
# logic:
# import raw end coordinate of modified (either elongated or shortened) exon
```

```
# if modified end coordinate at or downstream translation start (ts), event in CDS or in 3'UTR, otherwise (= upstream ts) event in 5'UTR

# in case of at or downstream ts: if modified end coordinate at or upstream translation end (te), event in CDS;

#  else (= downstream te), event in 3'UTR

  localization = 0

  raw_end_coordinate_modified_exon = 0

  # get raw end coordinate of modified exon:

  for entry in altd_list:

    if entry[0] == transcript_id:

      raw_end_coordinate_modified_exon = entry[1]

      break

  # comparisons between event coordinate and translation coordinates:

  if raw_end_coordinate_modified_exon >= raw_translation_coordinates[0]:

    # event in CDS or 3UTR

    if raw_end_coordinate_modified_exon <= raw_translation_coordinates[1]:

      # event in CDS

      print('* AltD in transcript isoform ', transcript_id, ' happened in CDS - AS event at ', raw_end_coordinate_modified_exon)

      localization = 2

    else:

      print('* AltD in transcript isoform ', transcript_id, ' happened in 3UTR - AS event at ', raw_end_coordinate_modified_exon)

      localization = 3

  else:

    # event in 5UTR

    print('* AltD in transcript isoform ', transcript_id, ' happened in 5UTR - AS event at ', raw_end_coordinate_modified_exon)

    localization = 1


  return localization


def get_exonskip_localization(transcript_id, exonskip_list, raw_translation_coordinates):

# determine whether exon skipping happened in 5'UTR, CDS, 3'UTR, 5'UTR+CDS, CDS+3'UTR, or 5'UTR+CDS+3'UTR

# transcript_id: transcript id of isoform to be examined

# exonskip_list_list: list of exonskip_list events, containing tuples of transcript id + tuple of coordinates needed for localization

# raw_translation_coordinates: Tuple containing raw start and end coordinate of translation

# Returns 0 for unsuccessful localization, 1 for 5'UTR, 2 for CDS, 3 for 3'UTR, 4 for 5'UTR+CDS, 5 for CDS+3'UTR, 6 for 5'UTR+CDS+3'UTR

  localization = 0

  raw_end_coordinate_exon_ahead_skipped_one = 0

  raw_start_coordinate_exon_after_skipped_one = 0
```

```
  raw_start_coordinate_skipped_exon = 0
  raw_end_coordinate_skipped_exon = 0

  # get raw start and end coordinate of skipped exon:
  # - find corresponding list entry in exonskip_list:
  for entry in exonskip_list:
    if entry[0] == transcript_id:
      raw_start_coordinate_skipped_exon = entry[1][0]
      raw_end_coordinate_skipped_exon = entry[1][1]
      raw_end_coordinate_exon_ahead_skipped_one = entry[1][2]
      raw_start_coordinate_exon_after_skipped_one = entry[1][3]
      break

  # comparisons
  if raw_translation_coordinates[1] <= raw_end_coordinate_exon_ahead_skipped_one:
    # event in 3'UTR
    print('* Exon skip in transcript isoform ', transcript_id, ' happened in 3UTR')
    localization = 3
    return localization
  if raw_translation_coordinates[0] >= raw_start_coordinate_exon_after_skipped_one:
    # event in 5'UTR
    print('* Exon skip in transcript isoform ', transcript_id, ' happened in 5UTR')
    localization = 1
    return localization
  if raw_translation_coordinates[0] <= raw_start_coordinate_skipped_exon and raw_translation_coordinates[1] >= raw_end_coordinate_skipped_exon:
    # event in CDS
    print('* Exon skip in transcript isoform ', transcript_id, ' happened in CDS')
    localization = 2
    return localization
  if raw_translation_coordinates[0] > raw_start_coordinate_skipped_exon and raw_translation_coordinates[0] < raw_start_coordinate_exon_after_skipped_one and raw_translation_coordinates[1] >= raw_end_coordinate_skipped_exon:
    # event in 5'UTR + CDS
    print('* Exon skip in transcript isoform ', transcript_id, ' happened in 5UTR and CDS')
    localization = 4
    return localization
  if raw_translation_coordinates[0] <= raw_start_coordinate_skipped_exon and raw_translation_coordinates[1] < raw_end_coordinate_skipped_exon and raw_translation_coordinates[1] > raw_end_coordinate_exon_ahead_skipped_one:
    # event in CDS + 3'UTR
```

```
    print('* Exon skip in transcript isoform ', transcript_id, ' happened in CDS and 3UTR')

    localization = 5

    return localization

   if raw_translation_coordinates[0] > raw_start_coordinate_skipped_exon and raw_translation_coordinates[1] < raw_end_coordinate_skipped_exon:

    # event in 5'UTR + CDS + 3'UTR

    print('* Exon skip in transcript isoform ', transcript_id, ' happened in 5UTR and CDS and 3UTR')

    localization = 6

    return localization


  return localization


def check_for_as_events(refdict2gene, transcriptModels, orf_data, nonrefdict2gene, ir_list, alta_list, altd_list, exonin_list, ii_list, unresolved_list, exonskip_list, as_event_list, as_duplicates):

  checked_for_as_events = False

  #print('check_for_as_events()')

  if refdict2gene != ():

    intronless_exons = get_intronless_exon_coordinates(refdict2gene[0], transcriptModels)

    #print(key, ': intron cleaned exons: ', intronless_exons)

    intronless_translation_data = get_intronless_translation_data(refdict2gene[0], transcriptModels, orf_data, intronless_exons)

    if intronless_translation_data[5] == False or intronless_translation_data[6] == False:

      # 1 or both boundary exons could not be determined

      print(intronless_translation_data[0], ': lacking exon data: no AS event investigation')

      return checked_for_as_events

     print(intronless_translation_data[0], ': intronless translation coordinates: (', intronless_translation_data[1], ', ', intronless_translation_data[2], ')')

      print(intronless_translation_data[0], ': boundary exons of translation: (', intronless_translation_data[3], ', ', intronless_translation_data[4], ')')

    raw_translation_coordinates = get_raw_translation_coordinates(refdict2gene[0], transcriptModels, orf_data, intronless_exons)

    print(refdict2gene[0], ': raw translation coordinates: (', raw_translation_coordinates[0], ', ', raw_translation_coordinates[1], ')')

    # check transcript isoforms:

    for key1, value1 in nonrefdict2gene.items():

      print(key1, ' is further isoform - comparison of exon boundaries required')

      as_event = detect_as_event(transcriptModels, refdict2gene[0], key1, ir_list, alta_list, altd_list, exonin_list, ii_list, unresolved_list, exonskip_list)

      # only count AS event, if it is new for the current gene (= deviating coordinate is different from previous deviating coordinates for a given as event type)

      # explains a discrepancy between a counted event and corresponding list length, which reflects all detected events

      if as_event[0] != 0:

        # only process event if it's not the same event at the same position
```

```
if as_event not in as_event_list:
  as_event_list.append(as_event)
  print('raw exon coordinate ', as_event[0], ' of transcript ', key1, ' deviates')
  # case differentiation according to AS event type:
  if as_event[1] == 1:
    # ir event
    localization = get_ir_localization(refdict2gene[0], key1, ir_list, intronless_translation_data)
    if localization == 1:
      ir_events['5UTR'] += 1
    if localization == 2:
      ir_events['CDS'] += 1
    if localization == 3:
      ir_events['3UTR'] += 1
    if localization == 0:
      ir_events['not localized'] += 1

  if as_event[1] == 2:
    # alta event
    localization = get_alta_localization(key1, alta_list, raw_translation_coordinates)
    if localization == 1:
      alta_events['5UTR'] += 1
    if localization == 2:
      alta_events['CDS'] += 1
    if localization == 3:
      alta_events['3UTR'] += 1
    if localization == 0:
      alta_events['not localized'] += 1

  if as_event[1] == 3:
    # altd event
    localization = get_altd_localization(key1, altd_list, raw_translation_coordinates)
    if localization == 1:
      altd_events['5UTR'] += 1
    if localization == 2:
      altd_events['CDS'] += 1
    if localization == 3:
      altd_events['3UTR'] += 1
    if localization == 0:
      altd_events['not localized'] += 1
```

```
if as_event[1] == 4:
  # exonin event
  localization = get_ir_localization(refdict2gene[0], key1, exonin_list, intronless_translation_data)
  if localization == 1:
    exonin_events['5UTR'] += 1
  if localization == 2:
    exonin_events['CDS'] += 1
  if localization == 3:
    exonin_events['3UTR'] += 1
  if localization == 0:
    exonin_events['not localized'] += 1

if as_event[1] == 5:
  # ii event
  localization = get_ii_localization(key1, ii_list, raw_translation_coordinates)
  if localization == 1:
    ii_events['5UTR'] += 1
  if localization == 2:
    ii_events['CDS'] += 1
  if localization == 3:
    ii_events['3UTR'] += 1
  if localization == 4:
    ii_events['5UTR+CDS'] += 1
  if localization == 5:
    ii_events['CDS+3UTR'] += 1
  if localization == 6:
    ii_events['5UTR+CDS+3UTR'] += 1
  if localization == 0:
    ii_events['not localized'] += 1

if as_event[1] == 6:
  # exonskip event
  localization = get_exonskip_localization(key1, exonskip_list, raw_translation_coordinates)
  if localization == 1:
    exonskip_events['5UTR'] += 1
  if localization == 2:
    exonskip_events['CDS'] += 1
  if localization == 3:
```

```
          exonskip_events['3UTR'] += 1
        if localization == 4:
          exonskip_events['5UTR+CDS'] += 1
        if localization == 5:
          exonskip_events['CDS+3UTR'] += 1
        if localization == 6:
          exonskip_events['5UTR+CDS+3UTR'] += 1
        if localization == 0:
          exonskip_events['not localized'] += 1

      if as_event[1] == 7:
        # mutually exclusive exon event
        mutexexon_events['not localized'] += 1

      as_event = (0, 0)
    else:
      # as_event in as_event_list - means: as event already detected for the currently investigated gene
      print('duplicate')
      if as_event[1] == 1:
        # ir event
        as_duplicates[0] += 1
      if as_event[1] == 2:
        # alta event
        as_duplicates[1] += 1
      if as_event[1] == 3:
        # altd event
        as_duplicates[2] += 1
      if as_event[1] == 4:
        # exonin event
        as_duplicates[3] += 1
      if as_event[1] == 5:
        # ii event
        as_duplicates[4] += 1
      if as_event[1] == 6:
        # exonskip event
        as_duplicates[5] += 1
checked_for_as_events = True
return checked_for_as_events
```

```
# import reference transcript data
orf_file = sys.argv[1]
orf_data = read_longest_orf_file(orf_file)

# import exon data of all transcripts
with open(sys.argv[2], 'rb') as fp:
    transcriptModels = pickle.load(fp)

txs = []
for t in orf_data:
  txs.append(t[0])
ref_tx = ''
not_investigated_genes = 0
ir_list = []
alta_list = []
altd_list = []
exonin_list = []
ii_list = []
unresolved_list = []
exonskip_list = []
as_event = ()
# examine same AS events for a gene only once; therefore the following list:
as_event_list = []
# count duplicates:
as_duplicates = [0,0,0,0,0,0]
intronless_exons = []
intronless_translation_data = ()
raw_translation_coordinates = ()
ir_events = {'5UTR':0, 'CDS':0, '3UTR':0, 'not localized':0}
alta_events = {'5UTR':0, 'CDS':0, '3UTR':0, 'not localized':0}
altd_events = {'5UTR':0, 'CDS':0, '3UTR':0, 'not localized':0}
exonin_events = {'5UTR':0, 'CDS':0, '3UTR':0, 'not localized':0}
ii_events = {'5UTR':0, 'CDS':0, '3UTR':0, '5UTR+CDS':0, 'CDS+3UTR':0, '5UTR+CDS+3UTR':0, 'not localized':0}
exonskip_events = {'5UTR':0, 'CDS':0, '3UTR':0, '5UTR+CDS':0, 'CDS+3UTR':0, '5UTR+CDS+3UTR':0, 'not localized':0}
mutexexon_events = {'not localized':0}
gene_name = ''
refdict2gene = ()
nonrefdict2gene = {}
tx_counter = 0
```

```
gene_counter = 0


for key, value in transcriptModels.items():
  tx_counter += 1
  if gene_name != key[0:9] and gene_name == '':
    # first gene - not yet examined
    gene_name = key[0:9]
    gene_counter += 1
  else:
    if gene_name != key[0:9]:
      # new gene, but not first one; previous gene completely examined
      if check_for_as_events(refdict2gene, transcriptModels, orf_data, nonrefdict2gene, ir_list, alta_list, altd_list, exonin_list, ii_list, unresolved_list, exonskip_list, as_event_list, as_duplicates) == False:
        not_investigated_genes += 1
      refdict2gene = ()
      nonrefdict2gene = {}
      gene_name = key[0:9]
      gene_counter += 1
      # reset as_event_list, because each gene is checked separately for event replicates
      as_event_list = []


  if key in txs:
    # reference transcript
    refdict2gene = (key, value)
  else:
    # transcript isoform
    nonrefdict2gene[key] = value


if gene_counter >= 1:
  if check_for_as_events(refdict2gene, transcriptModels, orf_data, nonrefdict2gene, ir_list, alta_list, altd_list, exonin_list, ii_list, unresolved_list, exonskip_list, as_event_list, as_duplicates) == False:
    not_investigated_genes += 1


print('examined genes: ', gene_counter)
print('proportion of examined genes for which no AS event check took place: ', not_investigated_genes)
print('examined transcripts: ', tx_counter)


print('')
print('Intron retention events: ', ir_events)
print('Alternative acceptor site events: ', alta_events)
```

```
print('Alternative donor site events: ', altd_events)

print('Exon insertion events: ', exonin_events)

print('Exon skipping events: ', exonskip_events)

print('Exitron events: ', ii_events)

print('Mutually exclusive exons events: ', mutexexon_events)

print('')

print('Duplicate intron retention events: ', as_duplicates[0])

print('Duplicate alternative acceptor site events: ', as_duplicates[1])

print('Duplicate alternative donor site events: ', as_duplicates[2])

print('Duplicate exon insertion events: ', as_duplicates[3])

print('Duplicate exitron events: ', as_duplicates[4])

print('Duplicate exon skipping events: ', as_duplicates[5])

print('')

print('Sum of intron retention events: ', len(ir_list))

print('Sum of alternative acceptor site events: ', len(alta_list))

print('Sum of alternative donor site events: ', len(altd_list))

print('Sum of exon insertion events: ', len(exonin_list))

print('Sum of exon skipping events: ', len(exonskip_list))

print('Sum of exitron events: ', len(ii_list))

print('Sum of unresolved events: ', len(unresolved_list))

print('')

print('all 5UTR events: ', ir_events['5UTR'] + alta_events['5UTR'] + altd_events['5UTR'] + exonin_events['5UTR'] + ii_events['5UTR'] + exonskip_events['5UTR'])

print('all CDS events: ', ir_events['CDS'] + alta_events['CDS'] + altd_events['CDS'] + exonin_events['CDS'] + ii_events['CDS'] + exonskip_events['CDS'])

print('all 3UTR events: ', ir_events['3UTR'] + alta_events['3UTR'] + altd_events['3UTR'] + exonin_events['3UTR'] + ii_events['3UTR'] + exonskip_events['3UTR'])

print('all 5UTR+CDS events: ', ii_events['5UTR+CDS'] + exonskip_events['5UTR+CDS'])

print('all CDS+3UTR events: ', ii_events['CDS+3UTR'] + exonskip_events['CDS+3UTR'])

print('all 5UTR+CDS+3UTR events: ', ii_events['5UTR+CDS+3UTR'] + exonskip_events['5UTR+CDS+3UTR'])

print('all not localized events: ', ir_events['not localized'] + alta_events['not localized'] + altd_events['not localized'] + exonin_events['not localized'] + ii_events['not localized'] + exonskip_events['not localized'] + mutexexon_events['not localized'])
```
